# Rhythmicity of Intestinal IgA Responses Confers Oscillatory Commensal Microbiota Mutualism

**DOI:** 10.1101/2021.10.11.463908

**Authors:** Hugo A. Penny, Rita G. Domingues, Maria Z. Krauss, Felipe Melo-Gonzalez, Suzanna Dickson, James Parkinson, Madeleine Hurry, Catherine Purse, Emna Jegham, Cristina Godinho-Silva, Miguel Rendas, Henrique Veiga-Fernandes, David Bechtold, Richard K. Grencis, Kai-Michael Toellner, Ari Waisman, Jonathan R. Swann, Julie E. Gibbs, Matthew R. Hepworth

## Abstract

Mutualistic interactions with the commensal microbiota are enforced through a range of immune responses that confer metabolic benefits for the host and ensure tissue health and homeostasis. Immunoglobulin (Ig)A responses directly determine the composition of commensal species that colonize the intestinal tract but require significant metabolic resources to fuel antibody production by tissue-resident plasma cells. Here we demonstrate IgA responses are subject to diurnal regulation by dietary-derived metabolic cues and a cell-intrinsic circadian clock. Rhythmicity in IgA secretion conferred oscillatory patterns on the commensal microbial community and its associated metabolic activity, resulting in changes to metabolite availability over the course of the circadian day. Our findings suggest circadian networks comprising intestinal IgA, the diet and the microbiota align to ensure metabolic health.

**One-Sentence Summary:** We demonstrate diurnal rhythms in intestinal IgA act to cross-regulate oscillations in the abundance of commensal microbes to foster mutualism.

## Introduction

Multiple mammalian species have evolved to maintain a finely balanced relationship with tissue-resident commensal bacteria that is mutually beneficial and critical for tissue homeostasis and the health of the organism. The commensal microbiota confers a multitude of mutualistic functions to mammalian hosts via the provision of complementary metabolic activity, immune regulation and colonization resistance that prevents outgrowth of pathogenic microbes (*1–4*). Healthy interactions between the host and commensal microbes are dynamically regulated and determined via a complex crosstalk between the microbiota - at both the individual species and community level - the intestinal immune system, metabolites and nutritional cues. Conversely, disruption of this network through changes in lifestyle, diet, infection or antibiotic use can precipitate the onset or progression of metabolic and inflammatory diseases (*2, 4–6*).

Immunoglobulin (Ig)A is a specialized antibody isotype that acts to regulate commensal bacteria community composition, tissue and niche residence and microbial gene expression (*7–9*). Within mucosal barrier tissues IgA is the dominant antibody isotype and is produced in a dimeric form bound by a *J chain* linker that facilitates it selective transport across the intact intestinal epithelium and secretion into the intestinal lumen (*7–9*). IgA is produced by tissue-resident plasma cells (IgA^+^ PC) predominantly found within the small intestine, and at higher quantities than any other antibody isotype at homeostasis – with estimates suggesting that several grams of IgA are produced per day in healthy humans (*10*). Plasma cells are terminally differentiated antibody-secreting lymphocytes of the B cell lineage, that dedicate the vast majority of their cellular capacity to expanded organelle function required to power antibody translation and secretion (*11, 12*). In line with this, the differentiation of a class-switched B cell to plasma cell is associated with a huge increase in cell-intrinsic cellular metabolism and nutrient transport (*12, 13*). Moreover, emerging evidence suggests changes in nutrition and diet can potently perturb IgA responses in the intestinal tract, with consequences for the microbiota and whole-body metabolism (*9, 14, 15*).

Taken together these findings highlight the significant metabolic requirements of maintaining mucosal antibody responses to reinforce homeostatic host-commensal bacteria interactions and mutualism. To minimize the energetic cost of such metabolically demanding biological axes many species have evolved dynamic regulatory mechanisms – most notably the regulation of key physiological processes through circadian rhythms. Circadian rhythmicity acts to align metabolically demanding processes with diurnal light cycles and feeding activity, thus temporally regulating biological activity during active periods - associated with feeding activity and potential immune challenges - or periods of rest. Mechanistically this is controlled by a hierarchically layered series of circadian clocks – most notably within the light-sensing suprachiasmatic nucleus of the brain – but also cell-intrinsic clocks present across a broad range of cell types in peripheral organs (*16–18*). At the molecular level this is controlled by a transcriptional feedback loop mediated by a series of core clock genes that counter-regulate their own transcription – thus imprinting rhythmicity, while also modulating a wider signature of genes to alter cell function (*17*). Indeed, it is now appreciated that many immune cells exhibit cell-intrinsic circadian-mediated control of cell migration and magnitude of effector functions (*19*). Furthermore, circadian misalignment - through altered dietary composition and feeding times, jet lag or shift work - has been associated with a number of metabolic and inflammatory diseases, suggesting a better understanding of circadian regulation of immunity will have therapeutic implications. However, the role of circadian rhythms in modulating intestinal immune crosstalk with the microbiota have only recently begun to be clarified (*20*), and remain incompletely understood.

Recent advances have also demonstrated diurnal oscillatory behavior within the composition and activity of the commensal microbiota itself, in part imprinted through immune pressures (*21–25*). Intriguingly it has also been proposed that bacteria may possess analogous circadian clock machinery (*26*), suggesting circadian rhythmicity and oscillatory biology may have evolved across species to bidirectionally regulate microbial mutualism with the mammalian host. Here, we report diurnal rhythmicity of the secretory IgA response and the IgA^+^ PC transcriptome and demonstrate roles for both the cell-intrinsic circadian clock machinery and cell-extrinsic feeding cues in aligning IgA responses. Critically, rhythmicity in IgA regulated oscillations in the composition and metabolic activity of the commensal microbiota, thus highlighting circadian regulation of the immune system and microbiota as a key determinant of microbial mutualism.

## RESULTS

### Intestinal IgA responses exhibit diurnal rhythmicity

We hypothesized that energetically demanding intestinal IgA responses may be subject to diurnal regulation. To test this, we assessed the levels of secretory IgA within the feces of a single cohort of C57BL/6 mice at five time points over a 24-hour day (Zeitgeber times; ZT0, 6, 12, 18, 0). The concentration of IgA detected in the feces was found to exhibit significant and marked variation over the day (Fig. 1A; p<0.0001 by JTK analysis and p<0.001 by One-Way ANOVA test; see methods), suggestive of diurnal oscillatory activity, and which remained evident after normalizing for minor variations in total protein content between samples (Fig. S1A). In contrast, we did not observe time of day differences in the frequency or cell numbers of tissue-resident IgA^+^ plasma cells (IgA^+^ PCs) within the small intestinal and colonic lamina propria (Fig. 1B–C and Fig. S1B-D), nor were diurnal oscillations observed in Peyer’s Patch-associated IgA class-switched germinal centre (GC) B cells (Fig. S1E-G). As IgA^+^ PC numbers and intestinal IgA secretion are most enriched in the small intestine (*27*), we next determined the intrinsic capacity of IgA^+^ PCs sort-purified at different times of the day to secrete IgA *ex vivo*. As expected, IgA^+^ PCs secreted high amounts of IgA into culture supernatants, unlike equal numbers of sort-purified IgD^+^ B cells or IgA^+^ B cells (Fig. S1H). Strikingly, IgA^+^ PCs from ZT0 secreted significantly higher IgA than equal numbers of cells sort-purified at ZT12 (Fig. 1D), suggesting the capacity of IgA^+^ PCs to secrete IgA - as opposed to the numbers of IgA^+^ PCs in the intestine - may determine diurnal rhythms in IgA secretion, as observed in the feces (Fig. 1A).

**Figure 1.**
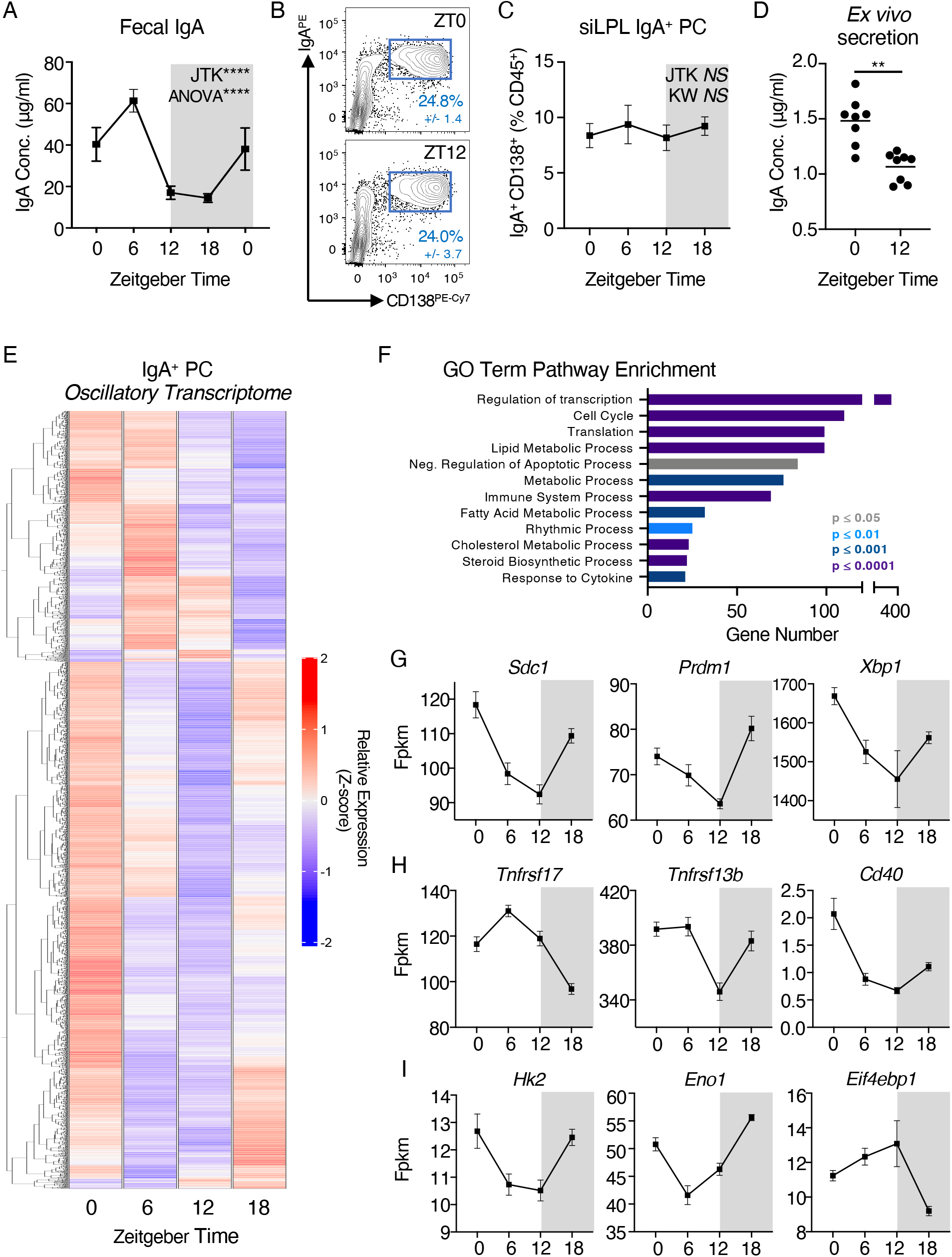
Mucosal antibody secretion and small intestinal IgA^+^ Plasma Cell activity exhibit diurnal rhythmicity. A) Serial fecal sampling of C57BL/6 mice at five 6 hour intervals over a circadian day (ZT 0, 6, 12, 18, 0), *n*=10 (pooled from two independent data sets). Data representative of at least 4 independent experiments. B) Exemplar flow plots of small intestinal CD138^+^ IgA^+^ plasma cells, pregated as Live CD45^+^CD3^−^CD5^−^NK1.1^−^MHCII^+/−^B220^−^IgD^−^, at ZT0 and ZT12 and C) Quantification of IgA^+^ plasma cell frequencies at ZT0, 6, 12 and 18. B+C *n*=5 and representative of three independent experiments. D) *Ex vivo* secretion of IgA by sort-purified IgA^+^ PC (from ZT0 and ZT12) cultured for 18 hours. Data pooled from two independent experiments, *n*=8. E) Heatmap of significantly oscillatory genes identified from bulk RNA Sequencing of sort-purified small intestinal IgA^+^ plasma cells taken at ZT0, 6, 12 and 18, z-score of average relative gene expression (fpkm) values of *n*=5 per timepoint. F) GO-Term pathway enrichment analysis on oscillatory gene signatures. Selected relative expression (fpkm) values for oscillatory gene signatures related to G) Plasma Cell function, survival and identitity, H) Extrinsic survival and antibody secretion signals, and I) Cellular Metabolism. All data shown as +/− SEM, * p< 0.05, ** p< 0.01, *** p< 0.001, **** p< 0.0001.

To further investigate the nature of diurnal regulation of IgA^+^ PC responses, we sort purified small-intestinal IgA^+^ PCs at ZT0, 6, 12 and 18 and performed bulk RNA-seq; of ~16,000 transcripts detected within our samples 2713 genes were found to exhibit highly significant time of day differences and oscillatory patterns after adjusting for false discovery rate (JTK analysis, BHQ <0.01), equivalent to ~16% of the observed transcriptome (Fig. 1E and Fig. S2A - top 50 differentially expressed genes). GO Term enrichment of highly oscillatory genes revealed enrichments in genes involved in Cell Cycle, Protein Translation, Metabolism and Rhythmic Process (Fig. 1F). Notably, oscillations were detected in the expression of key genes involved in IgA^+^ PC phenotype and transcriptional regulation (Fig. 1G), sensing of external activating signals and cell-cell crosstalk pathways known to influence antibody secretory activity (Fig. 1H), and metabolic activity and cholesterol biosynthesis pathways (Fig. 1I, Fig. S2B-G). Together, these findings suggest that IgA secretion and IgA^+^ PC-intrinsic transcriptional activity within the intestinal tract exhibits diurnal rhythmicity - and provoked the possibility of potential circadian entrainment.

### Cell-intrinsic circadian clock function is required for plasma cell transcriptional rhythmicity, but not rhythmic IgA secretion

Diurnal regulation of oscillatory transcriptional activity and function in both non-immune and immune cells is controlled in part by the cell-intrinsic “clock” - a transcriptional-translational feedback loop mediated by core clock proteins, including CLOCK, Bmal1 (encoded by *Arntl*), Rev-erbα (*Nr1d1*), Period (*Per1/2*) and Cryptochrome (*Cry1/2*). Notably, IgA^+^ PCs were found to have significant oscillations in the expression of *Arntl1*, *Nr1d1* and *Per2* by RNA Seq (Fig. S3A), which was independently validated via RT-PCR (Fig. 2A). As expected, expression of *Arntl* within IgA^+^ PCs was found to oscillate in anti-phase to *Nr1d1* and *Per2* over the circadian day, mirroring expression patterns in control liver tissue (Fig. S3B). In contrast, sort-purified naïve IgD^+^ B cells displayed no evidence of rhythmic expression of *Arntl* or *Per2*, although they surprisingly exhibited comparable oscillatory expression of *Nr1d1* (Fig. S3C). To determine the role of this cell-intrinsic circadian clock in regulating IgA secretion within the intestine, we generated conditional knockout mice in which Bmal1 was deleted within the B cell and PC lineages (*Mb1*^Cre^ x *Arntl*^fl/fl^). Efficient deletion of *Arntl* and disruption of associated clock gene transcription was confirmed in IgA^+^ PCs by RT-PCR (Fig. 2B and Fig. S3D).

**Figure 2.**
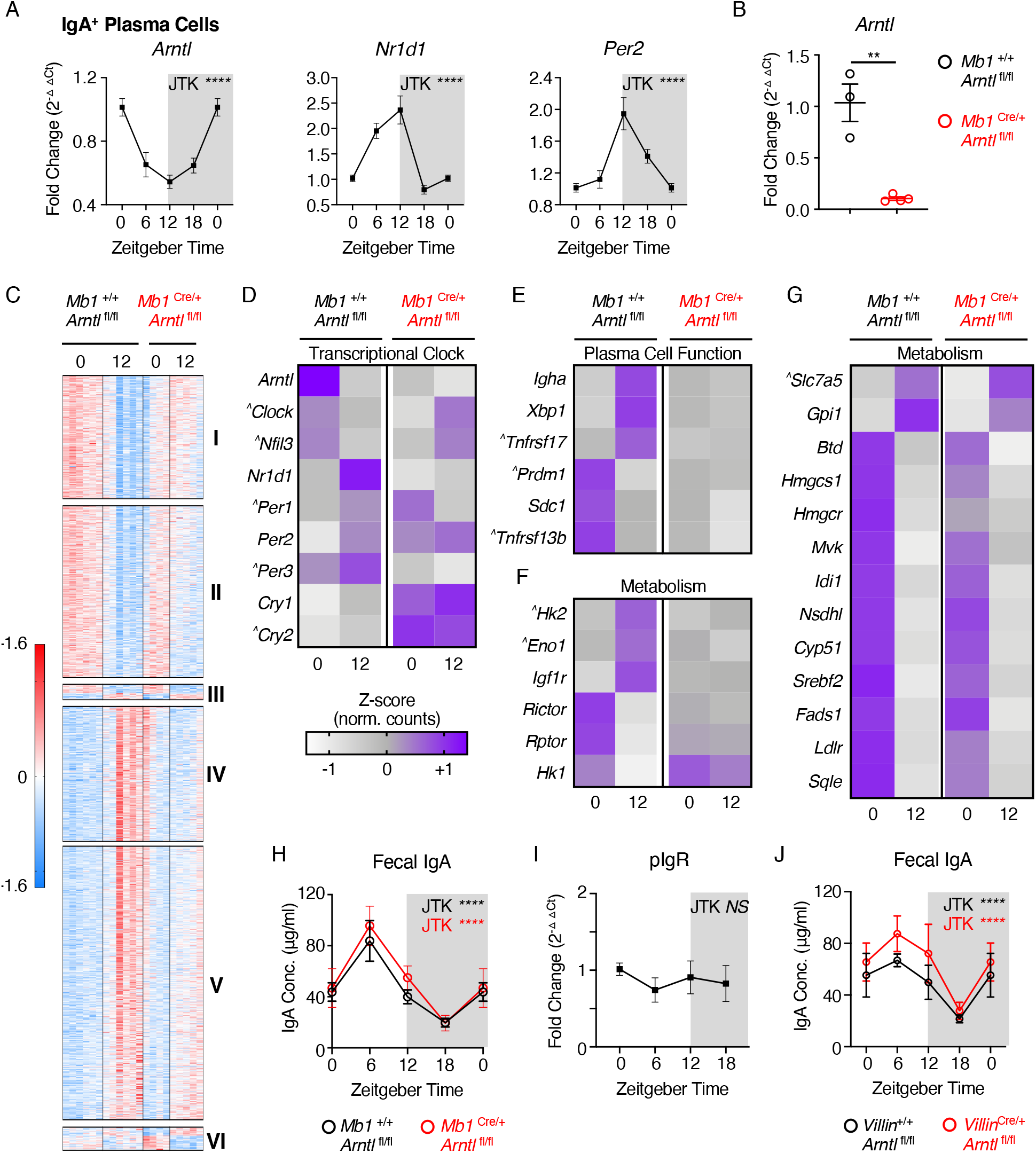
Rhythmic IgA^+^ Plasma Cell activity is in part dictated by the cell-intrinsic circadian clock. A) Relative expression of circadian clock genes in sort-purified small intestinal IgA^+^ PC at ZT 0, 6, 12 and 18 (ZT0 double plotted), determined by RT-PCR. *n*=10 (pooled from two independent experimental cohorts). Data representative of at least 3 independent experiments. B) RT-PCR validation of Arntl deletion in small intestinal IgA^+^ PC sort-purified from *Mb1*^Cre/+^ x *Arntl*^fl/fl^ mice in comparison to *Mb1*^+/+^ x *Arntl*^fl/fl^ littermate control animals, *n*=3-4 representativeof a single experiment. C) Heatmap comparison of differentially expressed genes identified by bulk RNA sequencing of sort-purified small intestinal IgA^+^ PC at ZT0 and 12, and found to significantly differ between ZT0 and ZT12 in control animals. Z-scores of relative gene expression (fpkm) values in individual animals of *n*=6 *Mb1*^+/+^ x *Arntl*^fl/fl^ mice and *n*=4-5 *Mb1*^Cre/+^ x *Arntl*^fl/fl^ mice per timepoint. Gene clusters: I+II (decrease in gene expression between ZT0 + ZT12 in controls, loss of suppression in *Mb1*^Cre/+^ x *Arntl*^fl/fl^ mice), III (time of day difference retained in both genotypes), IV+V (increase in gene expression between ZT0 + ZT12 in controls, loss of suppression in *Mb1*^Cre/+^ x *Arntl*^fl/fl^ mice) and VI (enhanced time of day difference in *Mb1*^Cre/+^ x *Arntl*^fl/fl^ mice). D-G) Average z-score values in IgA+ PC at ZT0 and ZT12 taken from *Mb1*^Cre/+^ x *Arntl*^fl/fl^ mice and *Mb1*^+/+^ x *Arntl*^fl/fl^ littermate control animals, representative of D) Circadian clock genes, E) Plasma Cell-associated genes, F+G) Metabolism-associated genes. ^ identifies genes where time of day differences either did not reach statistical significance in control animals in this analysis but were either previously identified in Figure 1 as oscillatory, or are directly related and relevant to the biological pathway described. H) Serial fecal sampling of *Mb1*^Cre/+^ x *Arntl*^fl/fl^ mice and *Mb1*^+/+^ x *Arntl*^fl/fl^ mice at four time points over a circadian day (ZT 0, 6, 12, 18; ZT0 double plotted), *n*=8-9 and pooled from two independent experimental cohorts. I) RT-PCR expression of *pIgR* relative to housekeeping gene in whole small intestinal tissue samples. J) Serial fecal sampling of *Villin*^Cre/+^ x *Arntl*^fl/fl^ mice and *Villin*^+/+^ x *Arntl*^fl/fl^ mice at four time points over a circadian day (ZT 0, 6, 12, 18; ZT0 double plotted), *n*=5 and representative of two independent experiments. All data shown as +/− SEM unless otherwise indicated, * p< 0.05, ** p< 0.01, *** p< 0.001, **** p< 0.0001.

To determine the role of IgA^+^ PC-intrinsic clock gene expression, we performed bulk RNA-seq on sort-purified small intestinal IgA^+^ PCs at ZT0 and ZT12 from *Mb1*^Cre^ x *Arntl*^fl/fl^ and wildtype littermate controls (Fig. 2C), and further confirmed severe disruption of time of day expression of the wider circadian clock gene family following deletion of *Arntl* (Fig. 2D). Analysis of differentially expressed genes revealed significant time of day-dependent signatures in control animals that were either lost (Cluster I and IV), suppressed (Clusters II and V), or retained (Cluster III) in the absence of a functional intrinsic clock (Fig. 2C). Additionally, we observed a time of day gene signature that was significantly enhanced in conditional knockout cells, when compared to controls (Cluster VI). Amongst these signatures we detected a loss of time of day differences in classical IgA^+^ PC-associated genes (Fig. 2E), as also identified in bulk RNA Seq analyses of wild type IgA^+^ PC over four time points (Fig. 1). In contrast, while a proportion of metabolism-associated genes displayed a clear loss of time of day differences in the absence of *Arntl* (Fig. 2F), others – including those involved in glycolysis, amino acid transport, the mevalonate pathway and cholesterol biosynthesis – retained time of day patterns (Fig. 2G), although in some cases the magnitude of this difference was altered or did not reach statistical significance.

Next we determined the impact of disrupted Bmal1-mediated regulation of PC transcription on rhythms in fecal IgA but unexpectedly found oscillations were retained (Fig. 2H), while IgA^+^ PC frequencies and numbers were unaffected by disruption of the cell-intrinsic circadian clock (Fig. S3E-F). As some time of day signatures in IgA^+^ PC transcription were only partly dependent on intrinsic *Arntl* expression and rhythmicity in IgA secretion was retained, we asked whether IgA secretion into the intestinal lumen could be subject to further circadian regulation at the tissue level. IgA produced by lamina propria-resident PCs requires active transport across the intestinal epithelium by the polymeric Ig receptor (pIgR). However, we failed to detect oscillatory expression of the *Pigr* gene in small intestinal tissue (Fig. 2I), while conditional deletion of *Arntl* in intestinal epithelial cells (*Villin*^Cre^ x *Arntl*^fl/fl^) also failed to perturb rhythmicity in fecal secretory IgA (Fig. 2J). Together these findings suggest that the IgA^+^ PC-intrinsic circadian clock is a major contributor to rhythmic transcriptional activity, but that rhythms in IgA secretion can persist in the absence of intrinsic clock function, indicating additional factors may entrain circadian function.

### Feeding-associated metabolic cues determine the magnitude and rhythmicity of intestinal IgA responses

While cell-intrinsic circadian clocks are important for driving oscillatory immune cell activity, additional exogenous signals can act to entrain these circadian rhythms - most notably feeding cues (*20, 28*). Moreover, emerging evidence suggests IgA responses are highly sensitive to changes in nutrition and diet (*9, 14, 15, 29*). To determine whether feeding-associated cues contribute to the entrainment of rhythms in IgA secretion, we utilized light-tight cabinets on reverse 12-hour light:dark schedules in combination with 6 hour periods of feeding restricted to either the dark phase (dark-fed) or light phase (light-fed) (Fig. 3A). Fecal sampling of animals maintained under these conditions at four time points (ZT0, 6, 12 and 18) revealed that dark-fed animals displayed oscillations in fecal IgA similar to that of *ad lib* fed mice (Fig. 3B, Fig. 1A), in line with the largely nocturnal feeding patterns of experimentally housed mice. Strikingly, restriction of food availability to a six-hour window during the light-phase led to a reversal in oscillatory IgA secretion (Fig. 3B and Fig. S4A) – indicating feeding cues act as a key entrainer of IgA secretion in the gastrointestinal tract. Notably, while IgA^+^ PC from dark-fed animals displayed cell-intrinsic time of day differences in clock gene expression comparable with *ad lib* fed mice, reversal of feeding also reversed clock gene expression patterns (Fig. 3C), which was mirrored in the liver (Fig. S4B). In line with our findings under *ad lib* conditions (Fig. 1 and Fig. S1), feeding cue-associated regulation of fecal IgA could not be attributed to alterations in IgA^+^ PC or IgA^+^ B cell frequencies in the intestinal tract and associated lymphoid structures (Fig. S4C+D).

**Figure 3.**
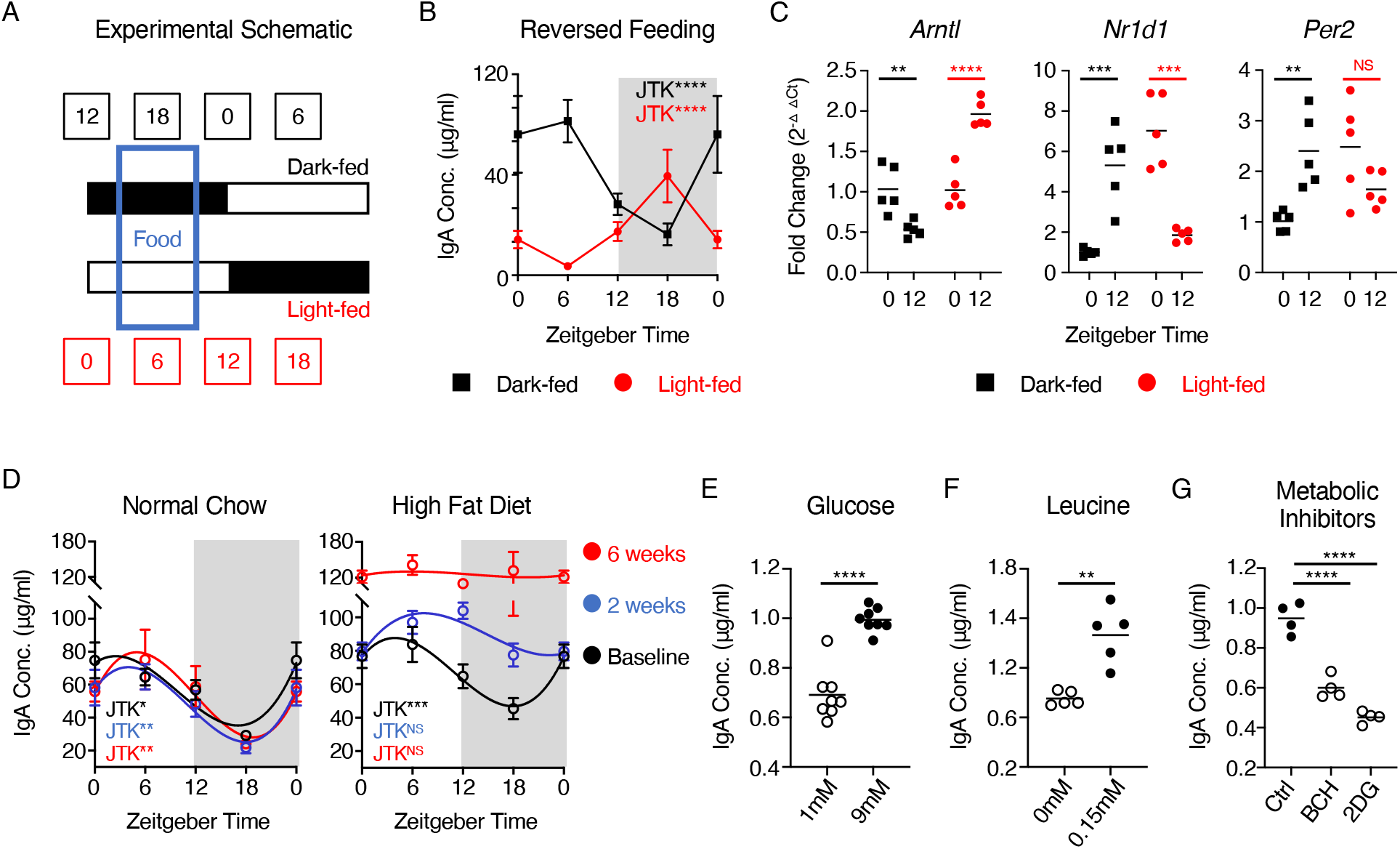
Oscillations in secretory IgA are aligned by feeding cues and cellular metabolic activity. A) Schematic of reversed feeding regimen, B) Serial fecal sampling of light-fed or dark-fed C57BL/6 mice at four 6 hour intervals over a circadian day (ZT 0, 6, 12, 18; ZT0 double plotted), *n*=9-10 (pooled from two independent experimental cohorts). Data representative of at least 4 independent experiments. C) RT-PCR analysis of circadian clock genes at ZT0 and ZT12 in sort-purified small intestinal IgA^+^ PC isolated from light-fed or dark-fed mice, *n*=5 per group, data representative of two independent experiments. D) Serial fecal sampling of C57BL/6 mice fed normal chow or high fat diet (HFD at five 6 hour intervals over a circadian day (ZT 0, 6, 12, 18; ZT0 double plotted), taken at baseline, two weeks or six weeks on the indicated diet, *n*=4-5 and data representative of at least 2 independent experiments. E-G) *Ex vivo* secretion of IgA by sort-purified small intestinal IgA^+^ PC cultured with differing concentrations of E) glucose (*n*= 8, representative of pooled data from two independent experiments) F) leucine (*n*= 5, representative of data from two independent experiments)or G) in the presence of metabolic inhibitors (*n*= 4, representative of data from three independent experiments). All data shown as +/− SEM unless otherwise indicated, * p< 0.05, ** p< 0.01, *** p< 0.001, **** p< 0.0001.

Together these findings suggested that feeding-associated cues, such as dietary-derived nutrients and metabolites, may act upstream to entrain cell-intrinsic clock genes - while also acting to regulate cell function through additional mechanisms independent of clock gene expression *per se*. As we observed time of day differences in a series of metabolic genes despite deletion of *Arntl* in IgA^+^ PC (Fig. 2G), we reasoned that feeding cues may further entrain IgA secretion via effects on plasma cell metabolic activity, in concert with clock gene driven regulation of transcription. We thus hypothesized that alterations in dietary nutritional content may perturb rhythms in IgA secretion. As proof of concept we fed mice normal chow or a commercial high fat diet (HFD), to establish a state of overnutrition, and assessed circadian rhythms in IgA secretion at baseline, 2 weeks or 6 weeks later. Animals fed HFD gained a moderate amount of weight over the 6-week period when compared to mice fed normal chow (Fig. S4E), and critically while post-prandial blood glucose was elevated in HFD mice after six weeks (Fig. S4F), no signs of metabolic disease or impaired glucose tolerance were observed at this time (Fig. S4G). In contrast, mice fed HFD for a prolonged period of 12 weeks began to exhibit elevated fasting glucose levels (Fig. S4G). Fecal IgA levels consistently exhibited circadian oscillations over a 24-hour period in animals fed normal chow and serially sampled at baseline, 2 weeks and 6 weeks (Fig. 3D). In contrast, while the HFD-fed group exhibited a comparable oscillation in fecal IgA at baseline, the same animals began to exhibit dysregulation of oscillatory IgA secretion following two weeks on HFD, and a complete loss of IgA rhythmicity after 6 weeks (Fig. 3D). Notably, the overall magnitude of IgA secretion was increased significantly in mice fed HFD for 6 weeks (Fig. 3D) suggesting excessive nutrition may nonetheless elevate IgA secretion in the intestinal tract over this time period.

Cell-intrinsic metabolic activity and nutrient availability have been demonstrated to be critical determinants of plasma cell survival, function and antibody secretory capacity (*12, 13, 30, 31*). In line with this concept, we found that small intestinal IgA^+^ PCs exhibited elevated metabolic activity when compared to either IgA^+^ B cells or IgD^+^ B cells derived from the Peyer’s patches (Fig. S4H-Q). Notably, IgA^+^ PCs exhibited markedly elevated uptake of the glucose analogue 2NDBG (Fig. S4H-I), expressed higher levels of the solute carrier chaperone protein CD98 – which functionally endowed cells with enhanced amino acid uptake capacity (Fig. S4J-N), and exhibited elevated intracellular lipid content (Fig. S4O-P). The heightened metabolic activity of IgA^+^ PCs was further reflected in extracellular flux assays (Fig. S4Q). Nonetheless, the *ex vivo* metabolic activity of IgA^+^ PCs did not significantly differ by time of day (Fig. S4R-U), suggesting that circadian rhythms in IgA^+^ PC function and IgA secretion were not dictated by diurnal changes in the metabolic capacity of plasma cells *per se*. Rather, we hypothesized that changes in nutrient availability - as a result of feeding activity - may act as a rate-limiting factor for antibody secretion by fueling IgA^+^ PC metabolism, and entraining rhythmicity in concert with the cell-intrinsic clock. In line with this hypothesis, the IgA secretory capacity of sort-purified PCs cultured *ex vivo* was found to be sensitive to the nutrient content of culture media, with an increase in glucose from subphysiological (1mM) to physiological (9mM) levels resulting in increased magnitude of IgA secretion (Fig. 3E). Similarly, IgA secretion from cultured PCs was sensitive to the presence of the amino acid leucine in the culture media (Fig. 3F), while pharmacological inhibition of either amino acid transport (BCH) or glycolysis (2DG) conversely reduced the magnitude of IgA secretion (Fig. 3G). Taken together these findings suggest that feeding-associated cues, through changes in nutrient availability, act to entrain and align oscillations in IgA production and the IgA^+^ PC transcriptional circadian clock in-part by fueling cell-intrinsic metabolic activity.

### Oscillatory IgA secretion partially imprints rhythms in the microbiota and modulates host-commensal mutualism

IgA is a canonical immune regulator of host-commensal microbe interactions and mutualism, and while a significant proportion of the microbiota is bound by secretory IgA the precise impact of IgA on the composition and mutualistic functions of the microbiota has remained incompletely understood. Conversely, emerging evidence suggests that the composition of the microbiota exhibits circadian rhythmicity which is in part dictated by host immune circuits (*20–23, 25, 32*). The dissection of the precise roles of IgA in regulating the commensal microbiota have been hindered by the observed generation of compensatory IgM responses in both *Igha*-knockout mice and IgA-deficient humans, which bind to a comparable repertoire of commensal bacteria (*27, 33–35*). Thus, to circumvent this issue and determine whether circadian oscillations in intestinal IgA impact upon the commensal microbiota we utilized IgMi mice, which lack the ability to class switch and secrete antibody yet retain a mature B cell compartment (*36–38*)(Fig. 4A). Thus, this model allowed us to study the microbiota in the absence of both secretory IgA and any other mucosal antibody isotypes transported into the intestinal lumen in the absence of IgA that may fully or partially compensate. As expected IgMi mice lacked detectable fecal IgA by ELISA when compared to littermate control animals (Ctrl) (Fig. S5A), and furthermore lacked IgA-bound bacteria as determined by flow cytometry (Fig. 4B, Fig. S5B). Next, we serially collected fecal samples from IgMi and littermate control animals over multiple circadian time points and performed 16S rRNA sequencing. In line with previous findings (*37*), we did not observe any dramatic changes in the global composition of fecal bacteria at the phylum or genus level when analyzing the microbiota of mice lacking IgA versus control animals, irrespective of circadian time (Fig. 4C; data shows average of combined timepoints). One notable exception was a clear reduction in the abundance of *Akkermansia* in IgMi mice (Fig. 4C and Fig. S5C-D), suggesting IgA binding may favor colonization of this mucosal-dwelling microbe. In contrast analysis by *Zeitgeber time* identified rhythms within the commensal microbiota, in line with previous reports (*21–23, 25, 32*). Consistent with these prior studies we were able to identify circadian rhythmicity in a number of bacteria genera including *Mucispirillum, Helicobacter*, *Peptococcaceae*, *Desulfovibrio* and *Bilophila* (Fig. 4D–E, H-I). In contrast, other major bacterial genera demonstrated no observable time of day differences (Fig. S5E). Critically, we identified a signature of rhythmic bacteria that lost circadian rhythmicity in the absence of IgA (Fig. 4D, H-I, Fig. S5F), although others retained or gained rhythmicity in IgMi mice (Fig. 4E–F). To determine whether the loss of bacterial rhythmicity in IgMi mice was preferentially observed amongst bacteria directly bound by IgA, we further performed IgA-Seq (Fig. 4G). IgA-Seq analysis revealed an enrichment (71%) of oscillatory microbes (*red*) amongst bacteria identified to be preferentially IgA bound in control animals. Notably, many of these bacteria also demonstrated a loss of rhythmicity (Fig. 4D and H), or changes in circadian phase (Fig. S5G) in the absence of mucosal antibody. Surprisingly, while we also observed a small subset of oscillatory bacteria that were preferentially enriched in the IgA negative fraction and unperturbed in IgMi mice (Fig. S5H), we also detected some bacteria that were not directly bound by IgA yet lost rhythmicity in IgMi mice (Fig. 4I) – suggesting circadian regulation of commensal communities may be complex and potentially subject to reciprocal interactions and competition for niche. Thus, we were able to identify oscillations in the abundance of a number of commensal bacteria that were dependent upon IgA, the secretion of which is itself regulated in a circadian manner.

**Figure 4.**
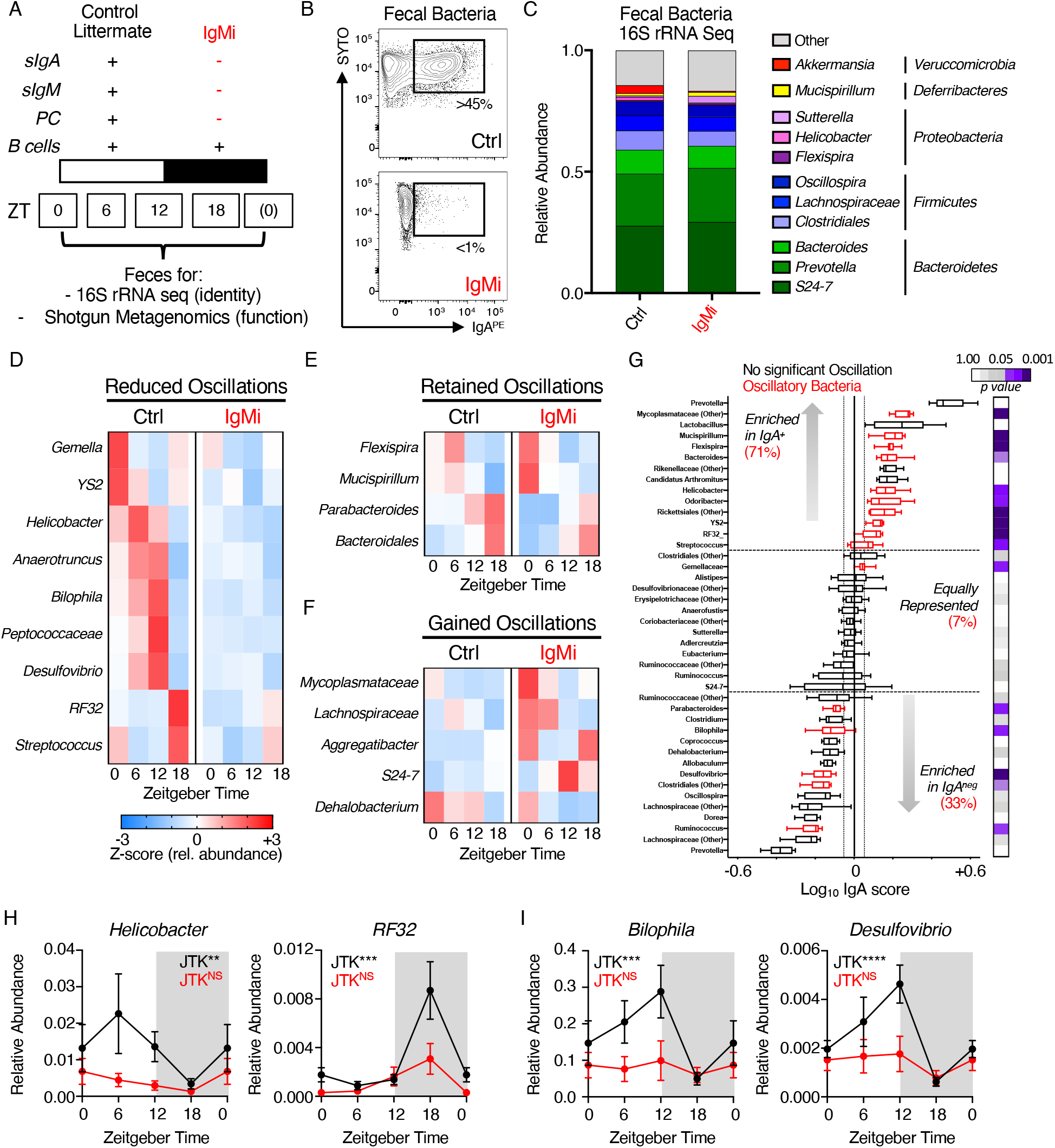
Rhythmic mucosal antibody production regulates circadian rhythmicity in the commensal microbiota. A) Summary of features of the IgMi mouse model. B) Representative measurement of IgA-binding to fecal bacteria in IgMi mice or littermate wild type control mice (Ctrl). C) Global analysis of average microbiota composition in Ctrl and IgMi animals elucidated by 16S rRNA Sequencing of fecal pellet-derived bacteria. D-F) Z-score heatmaps indicating average relative abundance of microbial genera in Ctrl and IgMi mice from serially sampled fecal bacteria taken at ZT0, 6, 12 and 18. G) IgA-Seq analysis of fecal bacteria isolated from Ctrl animals. Bacteria determined to exhibit oscillatory patterns in C-F are highlighted in red and the relative percent enrichment of oscillatory bacteria in IgA^+^ or negative fractions are indicated, *p* values for oscillatory analyses indicated in grey-purple. IgA enrichment indicate as log10 score. H+I) Individual data sets for selected bacteria identified as oscillatory in Ctrl animals and perturbed in IgMi mice, ZT0 data double plotted. All 16S rRNA sequencing and IgA Seq data representative of two independent experiments with *n*=4-5 animals per genotype, per ZT time point. All data shown as +/− SEM unless otherwise indicated, * p< 0.05, ** p< 0.01, *** p< 0.001, **** p< 0.0001.

While this provides evidence for a circadian role for IgA in regulating the composition of commensal bacteria, the consequences of this for the mutualistic functions of the microbiota and the mammalian host were unclear. Thus, we further performed shotgun metagenomics on serially sampled fecal bacteria from littermate control mice over five distinct time points (ZT0, ZT6, ZT12, ZT18 and a second ZT0, within the same 24-hour period). Analysis of functional GO-Terms in wild type control littermates predicted that a significant proportion of predicted bacterial functional pathways undergo circadian oscillation (Fig. 5A). Strikingly, IgMi mice exhibited a near-complete loss of highly oscillatory GO-Terms when compared to littermate controls (Fig. 5A+B). Many of the microbial GO-Terms that were found to be oscillatory in control animals and lost in IgMi mice related to metabolic processes, including *Glycolytic Process* and *Gluconeogenesis* (Fig. 5B+C, Fig. S6A+B), suggesting the presence of IgA may promote rhythmicity in microbial metabolism and liberation of nutrients from the diet. We also identified a small number of GO-Terms that indicated alterations in basic microbial biology, including several that in contrast were predicted to gain oscillations in the absence of IgA, including bacterial *Flagellum Assembly* and *Extrachromosomal Circular DNA* (Fig. S6C). Next, in order to determine whether changes in microbial function and metabolic activity altered nutrient availability within the intestine we performed metabolomics on fecal samples from IgMi mice and littermate controls. We observed evidence of time of day differences in the relative abundance of glucose in the feces over the course of a day, which were blunted in the absence of mucosal antibody (Fig. 5D), and to a lesser extent in short chain fatty acid availability (Fig. S7A), while availability of succinate exhibited comparable time of day differences regardless of the presence or absence of mucosal antibody (Fig. S7A). Despite changes in fecal metabolite levels, we confirmed that IgMi mice retained comparable circadian patterns in food intake (Fig. S7B), suggesting differences could not be attributed to changes in feeding behavior. To determine the potential impact of circadian changes in intestinal metabolite availability on the host we placed mice in metabolic cages (CLAMS) but found no evidence for major dysregulation of whole-body metabolism and energy usage (Fig. S7C+D). However, we observed perturbed time of day differences in circulating glucose in the blood of IgMi mice (Fig. 5E), which mirrored predicted microbial metabolic activity and glucose abundance in the feces, thus suggesting that circadian IgA regulation of microbial function may modulate time of day differences in metabolite availability and/or uptake by the host.

**Figure 5.**
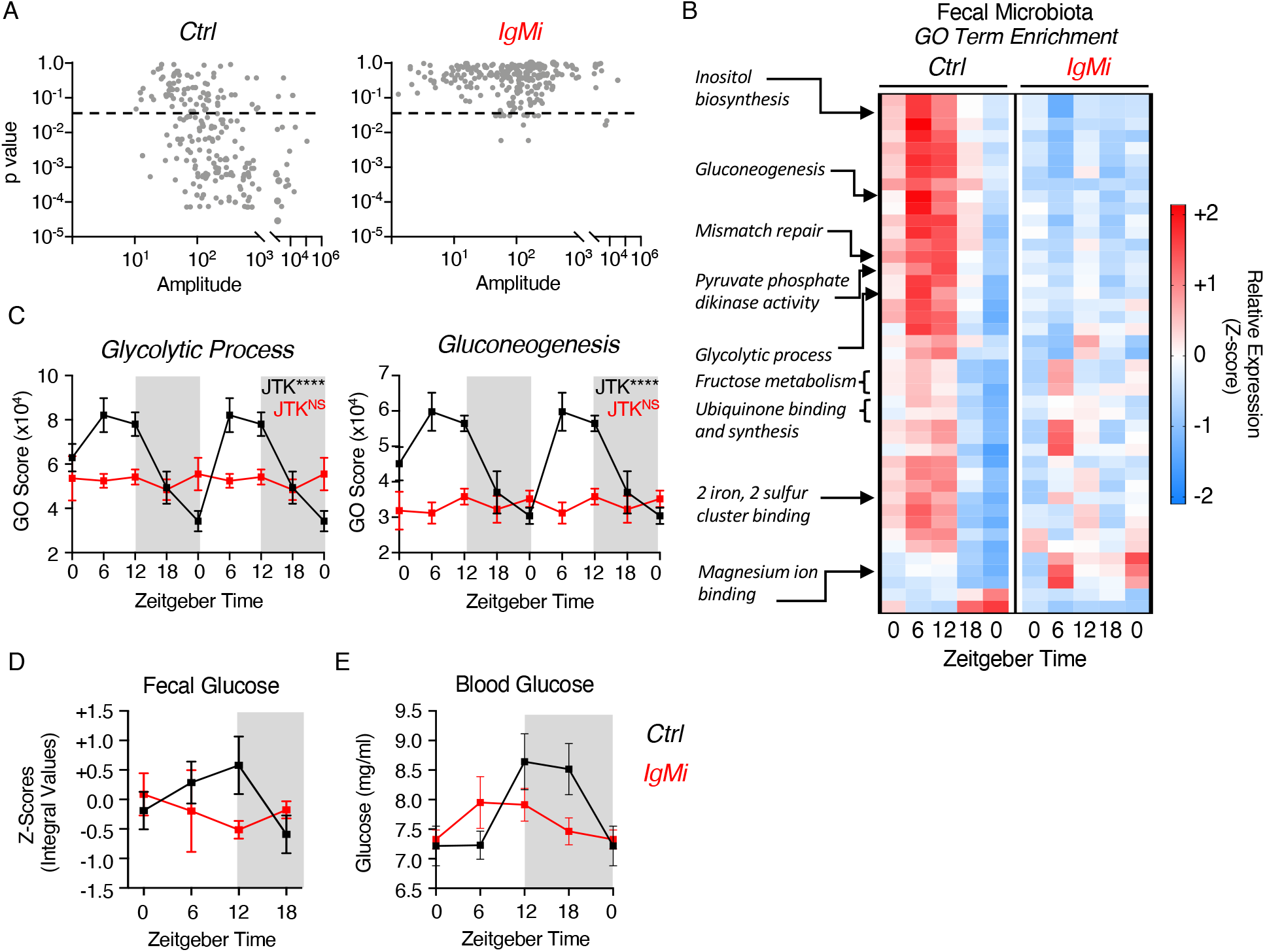
Mucosal antibody regulation of microbiome circadian rhythmicity modulates nutrient and metabolite availability and uptake. A) JTK analysis of GO Term pathway scores identified by shotgun metagenomics of serially sampled feces of Ctrl and IgMi mice over five 6 hour intervals over a circadian day (ZT 0, 6, 12, 18, 0), *n*=5 per group per timepoint, and representative of a single experiment. B) Z-score heatmap of average GO Term scores identified to be oscillatory in Ctrl mice and perturbed in IgMi mice, and C) select exemplar pathways double plotted. D) Glucose levels in serially sampled feces of Ctrl and IgMi mice over five 6 hour intervals over a circadian day (ZT 0, 6, 12, 18, 0), *n*=5 per group per timepoint, and representative of a single experiment. E) Glucose levels in serially sampled blood of Ctrl and IgMi mice over five 6 hour intervals over a circadian day (ZT 0, 6, 12, 18; ZT0 double plotted), *n*=8-12 per group per timepoint, representative of data pooled from three independent experiments. All data shown as +/− SEM unless otherwise indicated, * p< 0.05, ** p< 0.01, *** p< 0.001, **** p< 0.0001.

## DISCUSSION

The complex interplay between the microbiota, intestinal immune system and diet is increasingly understood to be central to a broad range of inflammatory, metabolic and systemic pathologies – and an increasing driver of morbidity and mortality in the industrialized world. In particular, an increased prevalence of high fat, low fibre diets and antibiotic use has been implicated in the onset and progression of obesity, allergy and chronic inflammation. Thus, understanding the consequences of interactions between commensal bacteria, intestinal immune responses and nutrition are key to disentangling the etiology and pathogenesis a broad range of diseases.

The constitutive regulation of host-microbiota interactions at mucosal barrier sites has the potential to be metabolically demanding. Here we provide evidence of circadian regulation of a major mucosal immune pathway central to host-microbiota crosstalk, which we hypothesize may have evolved to balance energetic cost with optimal orchestration of microbial mutualism. Specifically, we report diurnal secretion of IgA - in line with previous observations (*39, 40*) - and further define the precise cues and molecular mechanisms that align rhythms in mucosal antibody secretion as well the impact of this response on the microbiota. Our findings suggest a combination of cell-intrinsic circadian clocks and cell-extrinsic, feeding-associated nutritional cues entrain rhythms in IgA that modulate oscillations in the commensal microbiota and alter metabolite availability (Fig. S8). Intriguingly, the rhythmic regulation of glucose availability by IgA, and converse regulation of IgA oscillations by feeding-associated cues including glucose, suggests a bidirectional feedback loop may exist that tightly links IgA and commensal mutualism over the course of a day (Fig. S8).

Surprisingly while we observed IgA^+^ PC-intrinsic oscillations in canonical clock genes, and significant disruption of clock gene expression upon deletion of *Arntl*, rhythmicity in luminal secretory IgA was retained - suggesting Bmal1 is in part dispensable for oscillations in IgA secretion despite its clear role in regulating IgA^+^ PC transcription over time. This could be explained by findings that rhythmicity in both IgA secretion and clock gene expression was aligned by the time of feeding, in line with an emerging body of evidence that has demonstrated that rhythmic processes can be dictated in concert by both circadian and metabolic cues (*41, 42*). Nonetheless, we cannot rule out roles for circadian clock gene regulation independent of Bmal1. For example, other clock components have been reported to retain rhythmicity in the absence of Bmal1, while Rev-erbα has been attributed clock-independent roles as a transcription factor (*43*). Moreover, *Xbp1* – which is highly expressed by IgA^+^ PC – has recently been described to induce rhythmic gene expression independent of core clock genes (*44*).

In line with several previous studies a lack of IgA did not result in a marked dysbiosis *per se*, although we observed a loss of *Akkermansia* spp. in line with current understanding that in many cases IgA promotes host mutualism with mucosal-dwelling commensals (*45–47*). Strikingly however, we were able to recapitulate seminal observations made by other groups who reported diurnal oscillations in many of the same commensal microbes – including *Mucispirillum*, *Peptococcaceae* and *Streptococcaceae* spp (*21–23, 25, 32*). Critically, as in previous studies, these oscillations in bacterial constituents further manifested as time of day regulation of commensal function and broader microbial biology – most notably in pathways orchestrating nutrient metabolism, bacterial replication and pathogenicity (*21, 23, 25*). To date the role of IgA in impacting upon microbial *function* – as opposed to composition - has remained relatively poorly understood; here we demonstrate that lack of IgA secretion causes a loss in diurnal oscillations at the level of both composition and microbial activity, and consequently alters metabolite availability and host metabolic homeostasis.

Indeed, these data build upon previous findings in the field that suggest IgA binding has key roles in modulating bacterial gene expression, in addition to its more classical roles in regulating colonization and outgrowth. For example, we found that lack of IgA led to a gain in flagellum assembly over the circadian day, supporting findings that IgA binding can suppress bacterial flagellum expression (*48*). One notable observation was the circadian regulation of microbial pathways of glucose metabolism and glucose availability both within the intestine and circulation – adding to previous findings that IgA may be an important immune pathway in the regulation of glucose metabolism and risk of metabolic disease (*14*). Our data also build upon previous findings that indicate nutritional status and feeding-associated cues potently alter the magnitude of IgA secretion and microbiota-associated oscillations. Indeed, both long-term undernutrition or chronic overnutrition can alter the generation of IgA responses in the intestinal tract, suggesting a dynamic interplay between nutrition, circadian rhythms and mucosal antibody responses and host-commensal mutualism (*9, 14, 15, 20, 21, 23, 25, 28, 29*). More broadly, these findings suggest circadian IgA regulation of the microbiota may act to promote mutualism, metabolite availability and metabolic health, which together with recent advances (*49*), suggest IgA acts to determine host exposure to microbially-derived metabolites.

These findings complement and expand upon other recent studies that together suggest circadian regulation may be a common feature of tissue-resident intestinal immune cells that constitutively act to maintain healthy interactions with commensal bacteria (*24, 25, 43, 50–54*), and that immune pressure may partially imprint rhythmicity on the microbiota itself to confer mutualistic benefits for the host over the daily light:dark cycle, including ensuring energetic and metabolic efficiency aligned with feeding activity. An increasing body of evidence has begun to link lifestyles that disrupt circadian rhythmicity and microbial rhythms with the onset and progression of human inflammatory and metabolic diseases, including type 2 diabetes (*55*), and thus an increased understanding of circadian immune regulation will be critical to harness the full potential of the emerging field of circadian medicine (*56, 57*)

## MATERIALS AND METHODS

### Mice

Age and sex-matched C57BL/6 mice were purchased from Envigo laboratories. *Mb1*^Cre^ x *Arntl*^fl/fl^ were originally generated and provided by Kai-Michael Toellner (University of Birmingham), and IgMi mice were a kind gift from Ari Waisman (IMB Mainz). *Villin*^Cre^ x *Arntl*^fl/fl^ were maintained within the Centre for Biological Timing at the University of Manchester. All transgenic mouse experiments were performed using cohoused littermates and under specific pathogen free conditions with *ad libitum* feeding as 12h:12h light:dark cycle at the University of Manchester, United Kingdom, unless otherwise specified. In some cases, mice received irradiated High Fat Diet (Research Diets; D12492i; 60% Kcal from fat) *ad lib* for up to 12 weeks. Where indicated experimental cages were placed in controlled light-tight cabinets under opposing 12-hour light:dark cycles to facilitate investigation of circadian rhythms. In some experiments mice were placed in bespoke housing for the measurement of metabolic readouts and feeding as detailed below. All animal experiments were performed under Specific Pathogen Free (SPF) in single ventilated cages conditions and under license of the U.K. Home Office and under approved protocols at the University of Manchester.

#### Tissue Processing

Small intestinal lamina propria lymphocyte preparations were prepared by opening longitudinally and removing the Peyer’s patches, associated fat and luminal content by gently shaking in cold PBS. Epithelial cells and intra-epithelial lymphocytes were removed by shaking tissues in stripping buffer (1 mM EDTA, 1 mM DTT and 5% FCS) for two rounds of 20 min at 37°C. Lamina propria lymphocytes were isolated by digesting the remaining tissue in 1 mg/mL collagenase D (Roche) and 20 μg/mL DNase I (Sigma-Aldrich) for 45 min at 37°C. Liberated cells were then extracted by passing the tissue and supernatant over a 70μm nylon filter and centrifuged to isolate lamina propria lymphocytes. Isolated Peyer’s patches were processed by passing them through a 70μm nylon filter. In a small number of cases Peyer’s patches were retained during intestinal tissue digest to facilitate concurrent analysis of tissue-resident plasma cells and B cell subsets.

#### Flow Cytometry

Single cell preparations were stained with antibodies to the following markers: anti-CD3 (clone 145-2C11, eBioscience), anti-CD5 (clone 53-7.3, eBioscience), anti-B220 (clone RA3-6B2, eBioscience), anti-CD11b (clone M1/70, eBioscience), anti-MHCII (clone M5/114.15.2, eBioscience), anti-CD45 (clone 30-F11, Biolegend), anti-Fas (clone 15A7, eBioscience), anti-GL7 (clone GL7, Biolegend), anti-CD38 (clone 90, eBioscience), anti-CD98 (clone4F2, Biolegend) anti-CD19 (clone 1D3, BD), anti-IgA (clone mA-6E1, eBioscience), anti-IgD (clone 11-26c.2a, Biolegend), anti-CD138 (clone 281-2, Biolegend). Specific conjugates are indicated within Figures. Dead cells were excluded from analysis using the LIVE/DEAD Fixable Aqua Dead Cell Stain (Life Technologies). Samples were acquired using a BD Fortessa Cytometer, and analysed with FlowJo (TreeStar).

#### Bacterial flow cytometry

Feces were collected in Fast Prep lysing Matrix A tubes (MP Biomedicals), resuspended in 1ml of PBS per 100mg fecal material and incubated at 4°C for 20 min. Bacterial suspensions were resuspended in a final volume of 2 ml PBS and incubated at 4°C for 20 min. Samples were homogenized in a FastPrep-24 Tissue homogenizer (MP Biomedicals) for 30s. After homogenization, samples were centrifuged at 50 x *g* for 15 minutes at 4°C to remove debris and the bacteria-containing supernatant transferred through 70μm filters into a new tube. Bacteria were washed in FACs buffer (PBS, 2% FCS, 5mM EDTA) and pelleted at 8000 x *g* for 5 min. For flow cytometry, bacterial pellets were resuspended in 100μl FACs buffer containing SYTO 9 green fluorescent nucleic stain (Life Technologies) (10μM), incubated at 4°C for 15 minutes, and subsequently stained with 1μg/ml of an anti-mouse IgA-PE antibody (clone mA-6E1, eBioscience) for 30 min at 4°C. Samples were thoroughly washed and acquired on a BD Fortessa flow cytometer.

#### Cell-sorting and *ex vivo* culture assays

Kynurenine uptake was assessed as previously reported (*58*). Briefly, after surface antibody staining, 2×10^6^ cells were resuspended in 200μl warmed Hanks Balanced Salt Solution (HBSS; Sigma, UK), and 100μl of HBSS, or BCH (40mM, in HBSS), or leucine (20mM, in HBSS), was added to appropriate samples. Kynurenine (800μM, in HBSS) was then added and uptake subsequently stopped after 4 minutes by adding 125μl 4% PFA for 30min at room temperature in the dark. After fixation, cells were washed twice in HBSS and then resuspended in HBSS prior to acquisition on the flow cytometer. For assessment of 2-NBDG uptake *in vitro*, 1×10^6^ small intestinal cells were cultured in glucose-free DMEM medium (Agilent, USA) supplemented with 2mM L-glutamine and 100μM 2-NBDG (Thermo Fischer, USA) for 10 minutes at 37°C. Surface antibody staining of samples was then performed and acquisition of samples on the flow cytometer was undertaken within 2 hours. For assessment of lipid accumulation within cells *in vitro*, 1×10^6^ small intestinal cells were cultured in glucose-free DMEM medium (Agilent, USA) supplemented with 2mM L-glutamine and LipidTOX™ (Thermo Fischer, USA) for 30 minutes at 37°C. Cells were then washed, surface antibody staining of samples was then performed and acquisition of samples on the flow cytometer was undertaken within 2 hours.

#### ELISA

Mouse fecal IgA titers were measured using the Mouse IgA ELISA Quantitation Set (Bethyl Laboratories) following manufacturers’ instructions. Fecal samples were serially diluted and optimal dilutions and concentration were determined based via a standard curve. For core data sets an additional BCA assay (Pierce Coomassie Plus (Bradford) Assay Kit, Thermo Scientific) was performed on fecal extracts to measure total protein, and IgA concentrations normalized.

#### Metabolic inhibitor assays

Sort-purified IgA+ PCs isolated from the small intestinal lamina propria were incubated (10^4^ cells/ well) in either leucine free media (US Biological, USA) or glucose free media (Gibco, UK), with IL-6 (10ng/ml) (Peptrotech, USA) and BAFF (200ng/ml) (Biolegend, UK), supplemented with differing concentrations of leucine, or glucose (both Sigma, UK). To determine the effects of inhibiting nutrient uptake or metabolic signaling on IgA secretion, sort-purified IgA+ PCs isolated from the small intestinal lamina propria were cultured (10^4^ cells/ well) as above with or without the addition of metabolic inhibitors including pp242 (500nM), BCH (10mM) and 2-Deoxy-D-glucose (2DG) (1mM) (all Sigma, UK). Cells were incubated for 16 hours at 37°C, following which culture supernatants were removed and IgA concentrations determined by ELISA. Cell viability was determined under different culturing conditions, by either using a hemocytometer or flow cytometry.

#### Extracellular Flux Analysis

Extracellular flux analysis (Agilent, USA) was performed with replicates of 150,000 sort-purified IgA+ PCs isolated from small intestinal lamina propria or IgD+ B cells isolated from Peyer’s patches. Cells were adhered to each well of the Seahorse plate (Seahorse/Agilent, USA) using CellTak (Corning, USA). Cells were rested in Seahorse medium (glucose-free DMEM) at 37°C without CO2 for at least 30 minutes prior to the run. For the test, Seahorse XF medium was supplemented with 2mM of L-glutamine (Sigma, UK) and pH was adjusted to 7.35±0.05 (at 37°C). Glucose (10mM final concentration) (Fischer Scientific, USA), oligomycin (1μM final concentration (Sigma, UK) and 2-DG (100mM final concentration; Sigma), were added to individual ports to complete this assay.

#### Metabolic and physiological monitoring

To assess metabolic gas exchange, mice were individually housed in indirect calorimetry cages (CLAMS cages, Columbus instruments). Mice previously maintained on a controlled light-dark light cycle were acclimatized to the cages for two 24-hour cycles, and oxygen consumption and carbon dioxide production was recorded at 10-minute intervals for at least a further three consecutive 24h light-dark cycles. Respiratory exchange ratio (RER) was derived from these measurements (VCO2/VO2), as was energy expenditure. For measurement of food intake genotype-matched co-housed mice were placed in a Sable System for a full week on a controlled 24-hour light-dark cycle. Following a two-day acclimatization period, food intake was measured for at least three consecutives 24h light-dark cycles.

#### RT-PCR

Total RNA was purified using the RNeasy Micro Kit (Qiagen) and cDNA was prepared using the high capacity cDNA reverse transcription kit (Applied Biosystems). Real-time qPCR was performed with the real-time PCR StepOnePlus system (Applied Biosystems). TaqMan based assays (Applied Bio Systems) used the following primers and probes; *Gapdh* forward 5’ CAA TGT GTC CGT CGT CGA TCT 3’, Reverse 5’ GTC CTC AGT GTA GCC CAA GAT G 3’ and Probe 5’ CGT GCC GCC TGG AGA AAC CTG CC 3’; *Arntl* forward 5’ CCA AGA AAG TAT GGA CAC AGA CAA A 3’, Reverse 5’ GCA TTC TTG ATC CTT CCT TGG T 3’ and Probe 5’ TGA CCC TCA TGG AAG GTT AGA ATA TGC AGA A 3’; *Per2* forward 5’ GCC TTC AGA CTC ATG ATG ACA GA 3’, Reverse 5’ TTT GTG TGC GTC AGC TTT GG G 3’ and Probe 5’ ACT GCT CAC TAC TGC AGC CGC TCG T 3. *Nr1d1* was detected with a commercial Taqman probe assay (Mm00520708_m1; Applied Biosystems); Alternatively, LightCycler 480 SYBR Green I Master Mix (Roche) was used with the following primers; *pIgR* forward 5’ CTG GGG AAG AGG GAT CCA GA 3’ and reverse 5’ ACT CCC TTC ACA ACA GAG CG 3’, and *bactin* forward 5’ TCCTATGTGGGTGACGAG 3’ and *bactin* reverse 5’ CTCATTGTAGAAGGTGTGGTG 3’.

#### Bulk RNA sequencing

RNA was isolated from sort-purified cells, as above, and library preparation and bulk RNA sequencing was performed commercially with Novogene (UK) Company Ltd. Briefly, normalised RNA was used to generate libraries using NEB Next Ultra RNA library Prep Kit (Illumina). Indices were included to multiplex samples and mRNA was purified from total RNA using poly-T oligo-attached magnetic beads. After fragmentation, the first strand cDNA was synthesised using random hexamer primers followed by second strand cDNA synthesis. Following end repair, A-tailing, adaptor ligation and size section libraries were further amplified and purified and insert size validated on an Agilent 2100, and quantified using quantitative PCR (qPCR). Libraries were then sequenced on an Illumina NovaSeq 6000 S4 flowcell with PE150 according to results from library quality control and expected data volume. RNA Seq data are available via the GEO repository (Accession numbers: GSE175637, GSE175609).

#### 16S rRNA sequencing

Bacterial DNA from fecal bacteria was isolated using the PowerSoil DNA Isolation Kit (Qiagen, Netherlands) according to the manufacturer’s instructions. Pre-amplification of the V3V4 region of 16S rRNA was performed by PCR in triplicate using 2xKAPA HiFi Hot Start ReadyMix (Roche) using primer pairs containing adaptor sequences, as follows: 16S Amplicon PCR Forward Primer = 5′TCGT CGGCAGCGTCAGATGTGTATAAGAGACAGCCTACGGGNGGCWGCAG; 16S Amplicon PCR Reverse Primer = 5′ GTCTCGTGGGCTCGGAGATGTGTATAAGAGACAGGACT ACHVGGGTATCTAATCC. Following this, AMPure XP beads (Fisher Scientific) were used to purify the 16S V3V4 amplicon away from free primers and primer dimer species, according to the manufacturer’s protocol. Illumina sequencing adapters were then attached using the Nextera XT Index Kit (Illumina Inc, USA), according to the manufacturer’s instructions. DNA libraries were then further purified using AMPure XP beads. DNA libraries were then quantified, normalised and pooled together for 16S sequencing via the Illumina MiSeq platform (Illumina, USA) at the University of Manchester. IgA-Seq was performed as described previously (*36*), and sequencing performed at the University of Liverpool.

#### Shotgun Metagenomics

Shotgun metagenomics was performed commercially by CosmosID. Briefly, microbial DNA was extracted from fecal pellets and quantified using Qubit 4 fluorometer and HS Assay Kit (Thermofisher Scientific). DNA libraries were prepared using the Nextera XT DNA Library Preparation Kit and Nextera Index Kit (Illumina) following the manufacturer’s protocol with minor modifications. The standard protocol was used for a total DNA input of 1ng. Genomic DNA was fragmented using a proportional amount of Illumina Nextera XT fragmentation enzyme. Combinatory dual indexes were added to each sample followed by 12 cycles of PCR amplification. DNA libraries were then purified using AMPure magnetic beads (Beckman Coulter) and eluted in Qiagen EB buffer. DNA libraries were re-quantified and pooled together for sequencing via the Illimunia HiSeqX. Raw reads from metagenomics samples were analysed by CosmosID metagenomic software (CosmosID Inc., Rockville, MD, USA) to identify microbes to the strain level and a high-performance data mining k-mer algorithm was employed alongside highly curated dynamic comparator databases to rapidly disambiguate short reads into related genomes and genes.

#### Functional profiling of shotgun metagenomic data

Following initial QC, adapter trimming and preprocessing of metagenomic sequencing reads were performed using BBduk. The quality-controlled reads were then subjected to a translated search using Diamond against a comprehensive and non-redundant protein sequence database, UniRef 90. The mapping of metagenomic reads to gene sequences were weighted by mapping quality, coverage and gene sequence length to estimate community wide weighted gene family abundances. Gene families are then annotated to MetaCyc reactions (Metabolic Enzymes) to reconstruct and quantify MetaCyc metabolic pathways in the community. Furthermore, the UniRef_90 gene families were regrouped to GO terms to generate an overview of community function. To facilitate comparisons across multiple samples with different sequencing depths, the abundance values were normalized using Total-sum scaling (TSS) normalization to produce “Copies per million” units.

#### Bioinformatics

Where indicated bioinformatic analyses of data were performed via commercial platforms. For analysis of bulk RNA seq data differential gene expression analyses were performed in R (version 4.0.2) using RStudio Version 1.2.5033 (RStudio, Inc). Raw non-normalised counts were imported into R and subsequently analysed using the DESeq2 package (*59*), using the default pipeline. Genes with a total of fewer than ten counts across all samples were removed, and normalisation was calculated using the DESeq() function with default parameters for estimating size factors and dispersions. Differential expression was then calculated using the results() function with the default parameters. Genes with a significance value of less than 0.01 after correction for multiple comparisons using the Benjamini-Hochberg method were defined as “differentially expressed” and taken forward for further analysis. In some cases heatmaps were generated from normalised counts using the counts (normalised = TRUE) function followed by scaling and centring. Hierarchical clustering of genes was then computed using the ComplexHeatmap package (*60*). In other cases, clustering and normalised counts were then exported to excel and plotted in Graphpad Prism.

#### Metabolomics

The metabolic profiles of fecal samples were measured using ^1^H nuclear magnetic resonance (NMR) spectroscopy as previously described (*61*). Briefly, fecal samples (30 mg) were defrosted and combined with 600μL of water and zirconium beads (0.45 g). Samples were homogenized with a Precellys 24 instrument (45 s per cycle, speed 6500, 2 cycles) and spun at 14,000 *g* for 10 minutes. The supernatants (400μL) were combined with 250μL phosphate buffer (pH 7.4, 100% D_2_O, 3 mM NaN_3_, and 1 mM of 3-(trimethyl-silyl)-[2,2,3,3-^2^H4]-propionic acid [TSP] for the chemical shift reference at δ0.0) before centrifugation at 14,000 *g* for 10 minutes, and then transferred to 5 mm NMR tubes for analysis on a Bruker 700 MHz spectrometer equipped with a cryoprobe (Bruker Biospin, Karlsruhe, Germany) operating at 300 K. ^1^H NMR spectra were acquired for each sample using a standard one-dimensional pulse sequence using the first increment of the NOE pulse sequence for water suppression as previously described (*62*). Raw spectra were phased, baseline corrected and calibrated to TSP using Topspin 3.2 (Bruker Biospin) and then digitized in a Matlab environment (Version 2018; Mathworks Inc, USA) using in-house scripts. Redundant spectral regions (related to water and TSP resonance) were removed and the spectral data was manually aligned and normalized to the probabilistic quotient using in-house Matlab scripts. The peak integrals (relating to relative abundance) for metabolites of interest were calculated for each sample.

#### Statistical analyses

Statistical analysis of rhythmicity was calculated via JTK_Cycle analysis (*63*) of double plotted data sets using an established R pipeline. In some cases variations over time were additionally or alternatively analysed by Kruskal Wallis test or One Way ANOVA.

## ACKNOWLEDGEMENTS

We thank the Manchester Centre for Biological Timing, Gareth Howell, Mike Jackson, David Chapman in the University of Manchester Flow Cytometry facility, Andy Hayes and Claire Morrisroe in the University of Manchester Genomic Core, the University of Liverpool Centre for Genomics Research for 16S rRNA sequencing, and the University of Manchester BSF staff for support with animal husbandry and maintenance. We also acknowledge Suzanne Hodge, Giuseppe D’Agostino, Jenna Hunter, Devin Simpkins, Kathryn Gray and Edi Reshidi (all University of Manchester) for further support.

## FUNDING

Sir Henry Dale Fellowship jointly funded by the Wellcome Trust and the Royal Society, grant Number 105644/Z/14/Z (MRH)

BBSRC responsive mode, grant BB/T014482/1 (MRH)

Lister Institute of Preventative Medicine Prize (MRH).

Wellcome Trust Institutional Support Fund, grant 204796/Z/16/Z (MRH).

Wellcome Trust Centre for Cell Matrix Research, grant 203128/Z/16/Z (RKG).

Wellcome Trust 4ward North Clinical Fellowship, grant 203914/Z/16/Z (HAP).

EMBO long-term fellowship, grant ALTF 1209-2019 (RGD).

## AUTHOR CONTRIBUTIONS

Conceptualization: MRH, HAP, JG

Investigation: HAP, RGD, MZK, FMG, SD, MH, CP, EJ, CG-S, MR, DB, JS, MRH

Data Curation: HAP, RGD, JS, MRH

Formal Analysis: JP, HAP

Funding Acquisition: MRH, HAP, RGD.

Resources: DB, KMT, AW, HVF

Supervision: MRH, RKG, JG.

Writing – original draft: MRH

Writing – review and editing: HAP, RGD, MRH

## DATA AND MATERIALS AVAILABILITY

Data are included in the main text or supplementary materials. RNA Sequencing data sets will be made available via the aforementioned GEO accession numbers.

