## Supplemental Figures for "Rhythmicity of Intestinal IgA Responses Confers Oscillatory Commensal Microbiota Mutualism"

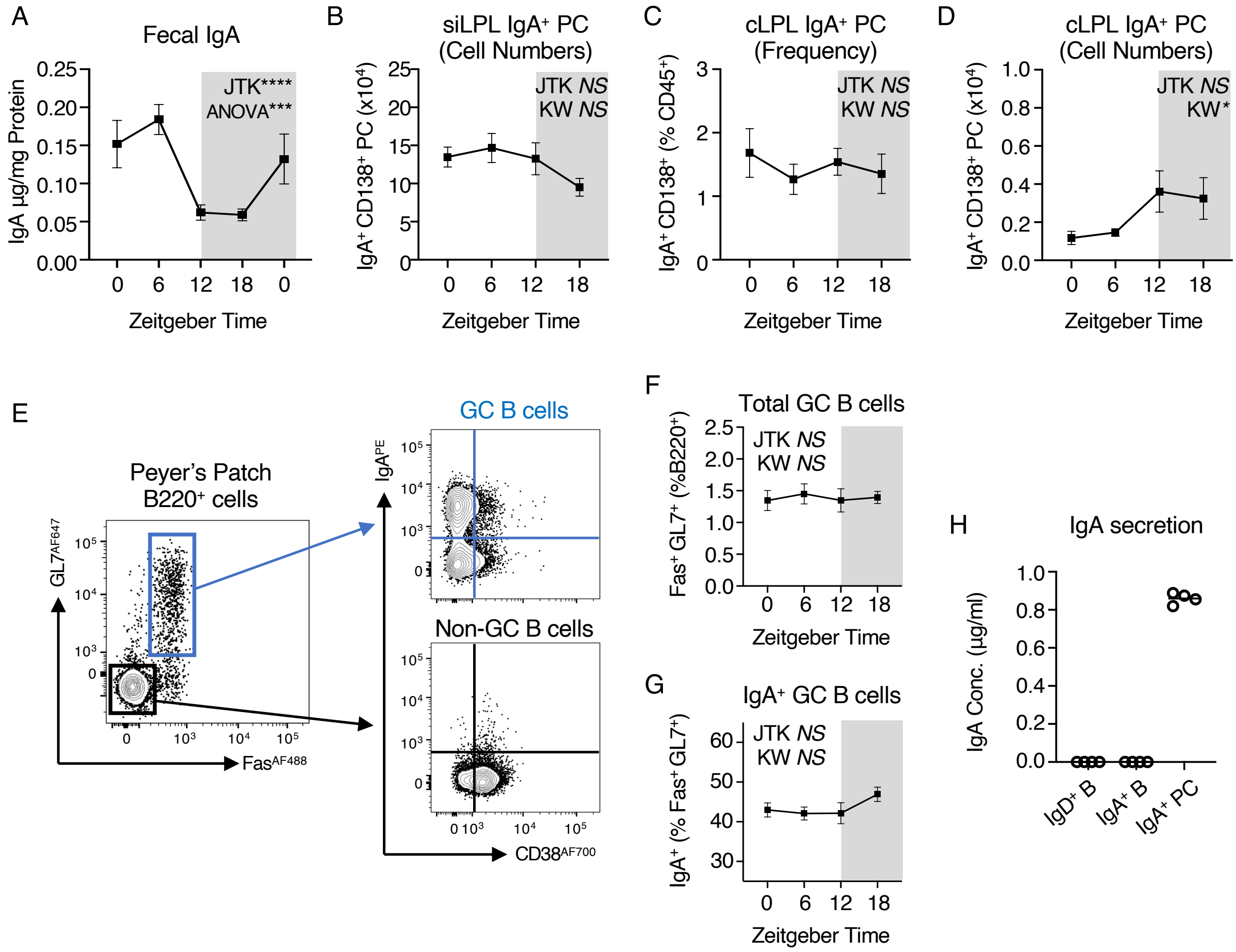


**Fig. S1. Rhythms in secretory IgA are not associated with diurnal oscillatons in intestinal plasma cell numbers or Peyer's patch responses.** A) IgA concentrations normalised to total protein concentrations of feces, serially sampled from C57BL/6 mice at five 6 hour intervals over a circadian day (ZT 0, 6, 12, 18, 0), *n*=10 (pooled from two independent experimental cohorts). Data representative of at least 2 independent experiments. B-D) Quantification of B) small intestinal lamina propria $IgA^+$ PC cell numbers, C) colon lamina propria frequencies (of live $CD45^+$ cells) and D) cell numbers of $IgA^+$ PC. B-D *n*=10 and representative of pooled data from two independent experiments. E) Representative flow plots indicating gating strategy for identification of Germinal Center (GC) B cells and IgA class-switched GC B cells in the Peyer's Patches. Cells pre-gated as Live $CD45^+$CD3-CD5-NK1.1-CD11b-$MHCII^+B220^+$. F) Frequencies of total GC B cells within Peyer's Patch B cell compartment and G) $IgA^+$ cells amongst GC B cells, from C57BL/6 mice sacrificed at four 6 hour intervals over a circadian day (ZT 0, 6, 12, 18), *n*=10 (pooled from two independent experimental cohorts). H) Secretion of IgA into culture media after overnight culture of equal numbers of sort-purified small intetinal $IgA^+$ PC, Peyer's Patch-derived $IgA^+$ B cells or $IgD^+$ B cells. All data shown as +/- SEM unless otherwise indicated, * $p< 0.05$, ** $p< 0.01$, *** $p< 0.001$, **** $p< 0.0001$.

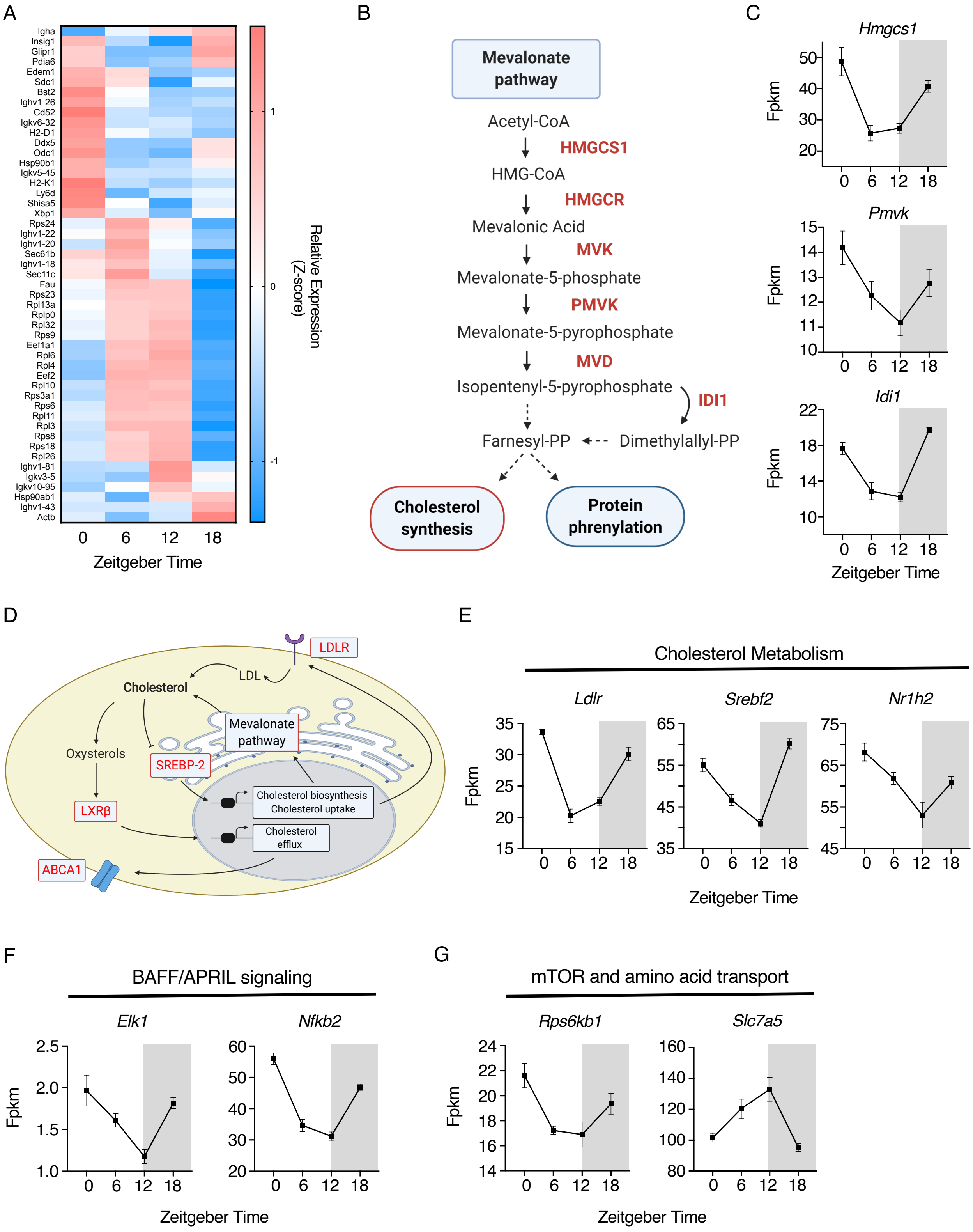


**Fig. S2. Diurnal rhythmicity in IgA$^+$ Plasma Cell function is associated with oscillatory activity in metabolic pathways and sensing of survival signals.** A) Z-score heatmap of top 50 significantly oscillatory genes identified in small intestinal IgA$^+$ PC sort-purified at ZT0, 6, 12 and 18. Average values of *n*=5 per timepoint. B) Enrichment of sigificantly oscillatory genes (highlighted in red) in the Mevalonate pathway. C) Relative expression (fpkm) of selected genes from B. D) Representative image indicating significantly oscillatory genes associated with cholersterol biosynthesis and metabolism and E) relative expression (fpkm) of selected genes from D. F) Relative expression (fpkm) of additional genes associated with pathways highlighed in Figure 1H. G) Relative expression (fpkm) of additional genes associated with pathways higlighed in Figure 1I.

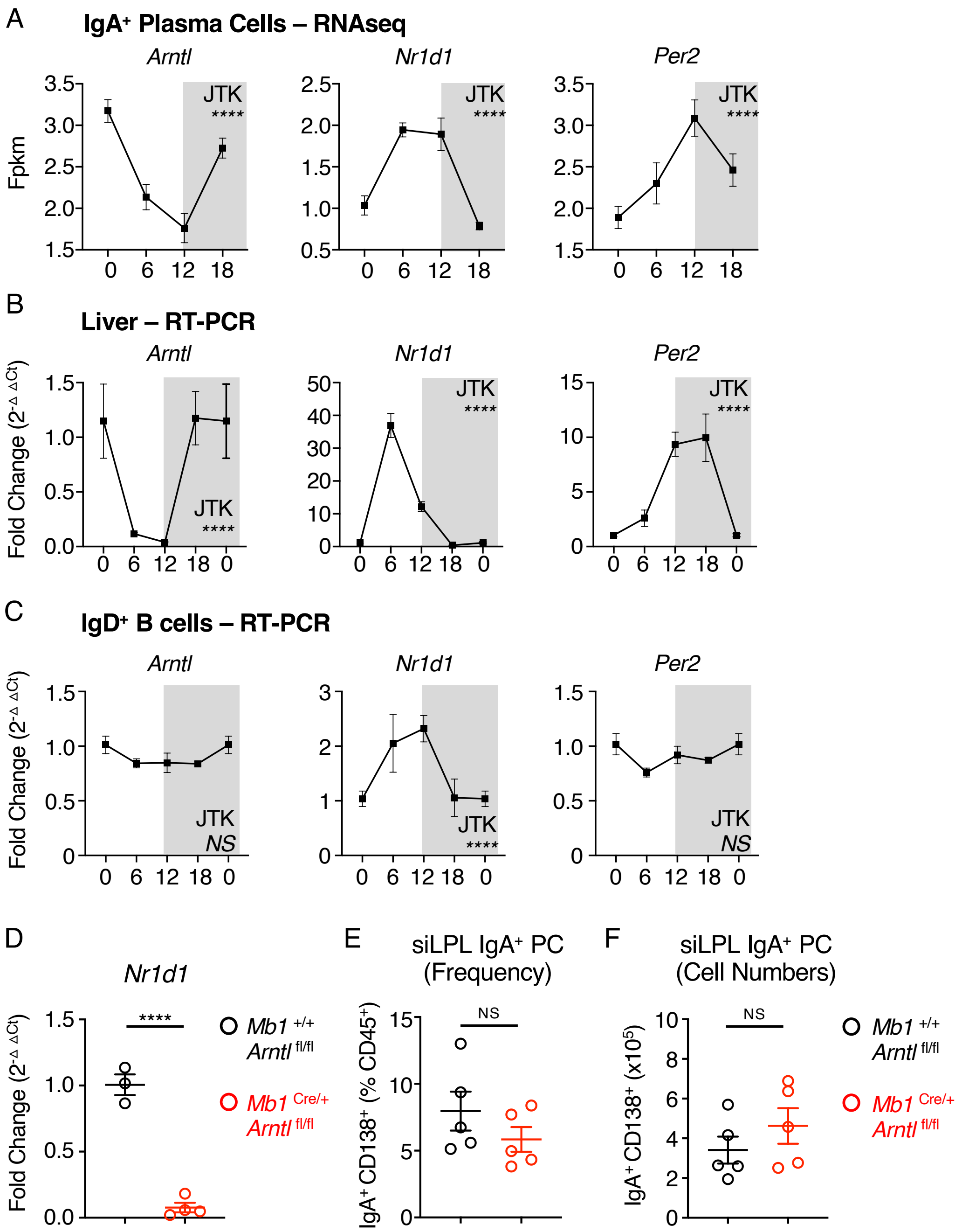


**Fig. S3. Circadian clock gene expression and validation of a conditional *Arntl*- knockout mouse model.** A) Relative expression (fpkm) of circadian clock genes as identified by bulk RNA seq in Figure 1. B) RT-PCR validaton of steady state circadian clock phase in whole liver tissue samples from mice culled at ZT0, 6, 12 and 18 (ZT0 double plotted). $n$=3-4 and representative of two independent experiments. C) RT-PCR validation of circadian clock genes in B220$^{+}$ IgD$^{+}$ B cells isolated from Peyer's Patches and small intestinal lamina propria of C57BL/6 mice culled at ZT0, 6, 12 and 18 (ZT0 double plotted). $n$=10 and representative of pooled data taken from two independent experiments. D) *Nr1d1* expression in small intestinal IgA$^{+}$ PC sort-purified from *Mb1*$^{\text{Cre/+}}$ x *Arntl*$^{\text{fl/fl}}$ mice in comparison to *Mb1*$^{+/+}$ x *Arntl*$^{\text{fl/fl}}$

littermate control animals, *n*=3-4 mice per group. E) Frequency and F) numbers of small intestinal $IgA^+$ PC in comparison to $Mb1^{+/+}$ x $Arntl^{fl/fl}$ littermate control animals, *n*=5 representative of two independent experiments. All data shown as +/- SEM unless otherwise indicated, * $p< 0.05$, ** $p< 0.01$, *** $p< 0.001$, **** $p< 0.0001$.

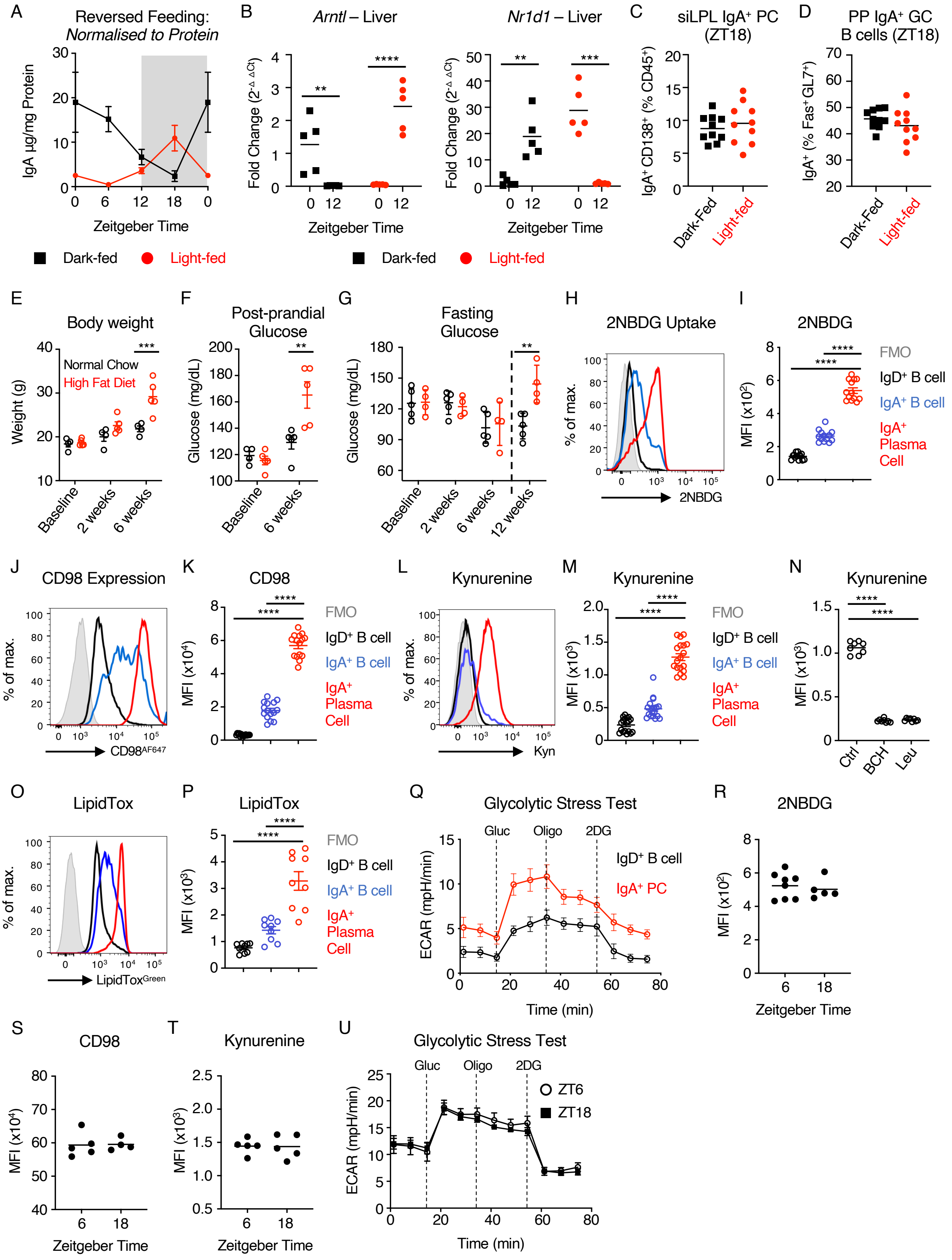


**Fig. S4. IgA$^{+}$ plasma cells exhibit elevated metabolic activity and comparable capacity to utilize metabolic substrates over the course of a circadian day.** A) IgA concentrations normalised to total protein concentrations of feces, serially sampled from light-fed and dark-fed C57BL/6 mice at four 6 hour intervals over a circadian day (ZT 0, 6, 12, 18; ZT0 double plotted), *n*=9-10 (pooled from two independent experimental cohorts). Data representative of at least three independent experiments. B) Validation of circadian clock dysregulation by reverse feeding in whole liver tissue. RT-PCR analysis at ZT0 and ZT12 in samples taken from light-fed or dark-fed mice, *n*=5 per group, data representative of two independent experiments. Quantification of C) small intestinal $IgA^+$ PC frequencies (% of total $CD45^+$) or D) Peyer's Patch $IgA^+$ GC B cells (% of GC B cell compartment) in light-fed or dark-fed mice, *n*=10 per group, data pooled from two independent experiments. E) Body weight of mice, F) post-prandial blood glucose (*ad lib* fed, sampled at ZT1) and G) fasting glucose (following 8 hour fast) in normal chow and high fat diet fed mice as in Figure 3. H-P) Assays to determine metabolic activity of $IgA^+$ PC, $IgA^+$ B cells and $IgD^+$ B cells from small intestinal lamina propria and Peyer's patches. Data pooled from two independent experiments. H+I) Uptake of 2-NBDG, J+K) CD98 expression, L+M) Kynurenine uptake and N) Kynurenine uptake in the presence of the amino acid inhibitor BCH, or exogenous cold Leucine, O+P) LipidTox staining. Q) Glycolytic stress test of sort-purified small intestinal-derived IgA+ PC and Peyer's patch-derived $IgD^+$ B cells by extracellular flux analysis. Data representative of two independent experiments. Comparison of R) 2-NBDG uptake, S) CD98 expression, T) Kynurenine uptake and U) glycolytic capacity in $IgA^+$ PC at ZT 6 and ZT18. All data shown as +/- SEM unless otherwise indicated, * $p< 0.05$, ** $p< 0.01$, *** $p< 0.001$, **** $p< 0.0001$.

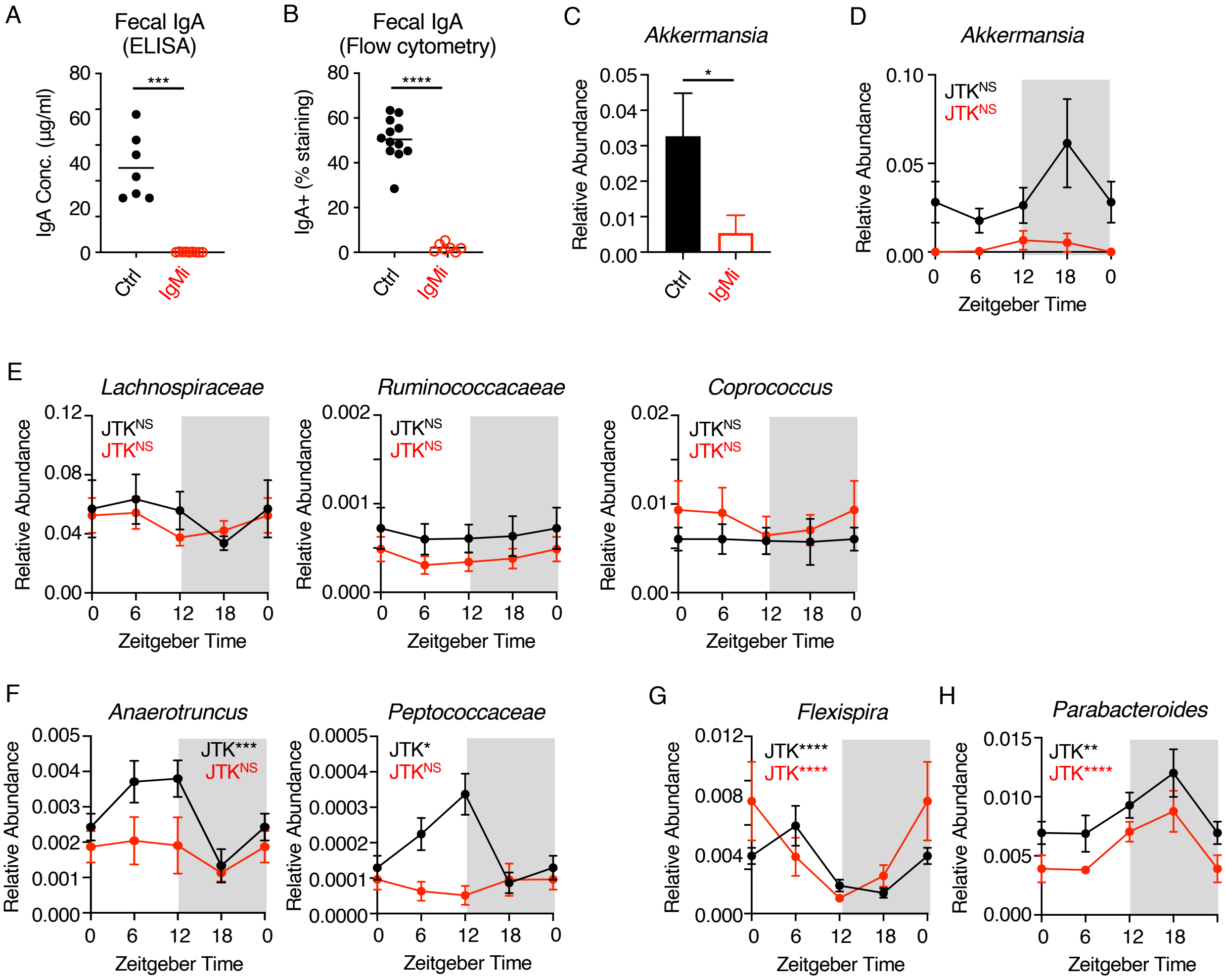


**Fig. S5. IgMi mouse validation and circadian rhythms in microbial abundance.** A) Fecal IgA in IgMi versus control littermate animals. B) IgA binding of fecal commensal baceria in IgMi and control littermates assessed by flow cytometry. C+D) Relative abundance of *Akkermansia* spp. in IgMi and Ctrl mice assessed via C) global analysis or D) ZT analysis. E) Selected commensal bacteria exhibiting no significant oscillatory behaviour in IgMi or Ctrl animals. F) Selected commensal bacteria exhibiting significant oscillatory behaviour in Ctrl but not IgMi animals but not detected in IgA Seq analysis, related to Figure 4D and 4G. G) Selected commensal bacteria exhibiting a retained but significant phase shift in oscillatory behaviour between Ctrl and IgMi animals, related to Figure 4E, and H) Selected commensal bacteria exhibiting a significant oscillatory behaviour that is retained in both Ctrl and IgMi animals, related to Figure 4E. All data shown as +/- SEM unless otherwise indicated, * $p < 0.05$, ** $p < 0.01$, *** $p < 0.001$, **** $p < 0.0001$.

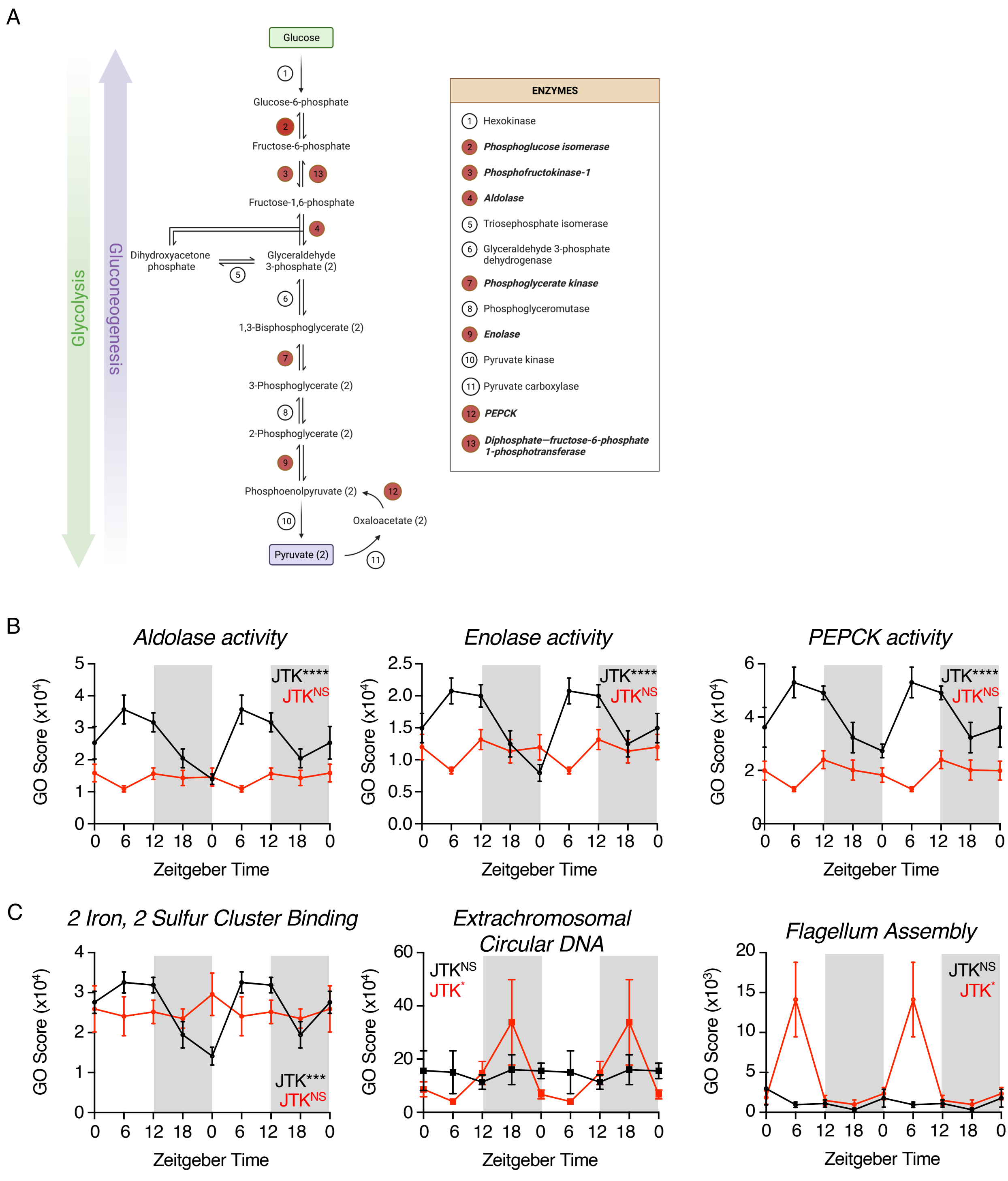


**Fig. S6. Alterations in predicted microbial gene pathways in the absence of mucosal antibody.** A) Schematic of GO Term pathways related to Figure 5C and glucose metabolism. Steps highlighted in red indicate significant oscillatory steps and/or enzymatic activity. B) Selected individual data for pathways related to A. C) Selected individual data of microbial GO Term pathways either lost or gained in IgMi mice in comparison to Control animals. All data shown as +/- SEM unless otherwise indicated, * $p < 0.05$, ** $p < 0.01$, *** $p < 0.001$, **** $p < 0.0001$.

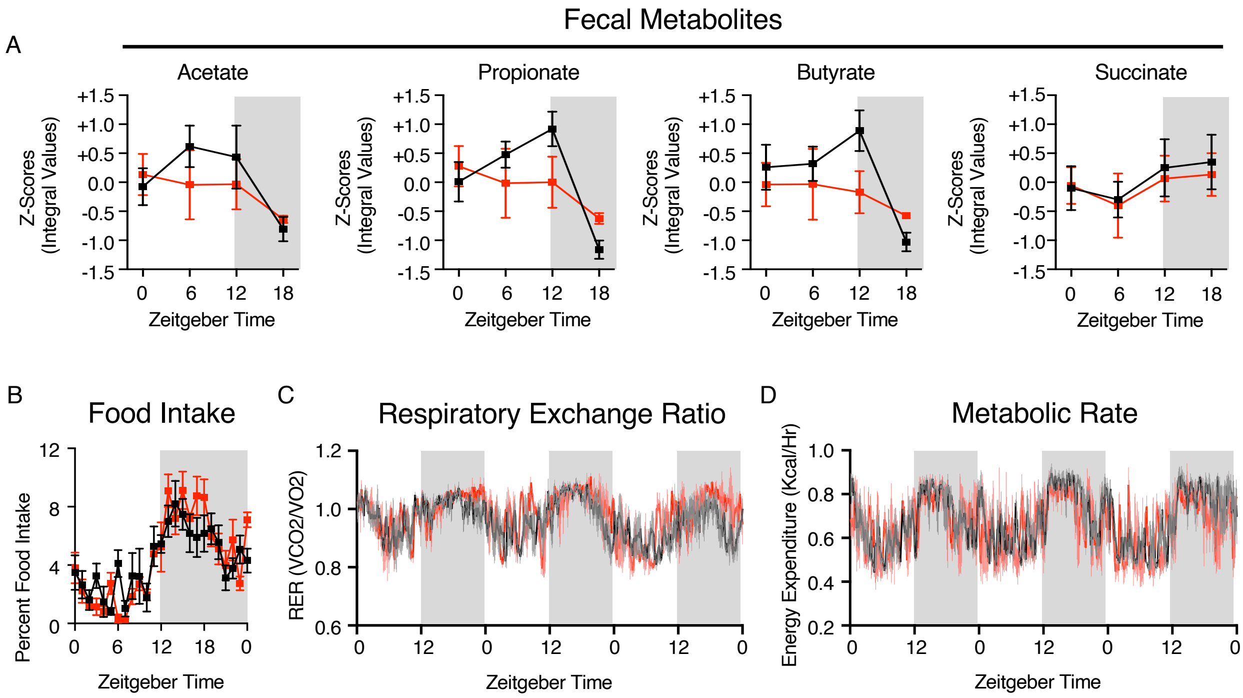


**Fig. S7. Dysregulation of circadian metabolites in the absence of mucosal antibody.** A) Additional fecal-associated metabolites over four time points in IgMi and Ctrl animals. Related to Figure 5D. B) Measurement of food intake at 1 hour intervals over three consecutive days in IgMi and Ctrl animals, *n*=4 per group and representative of a single experiment. C) Measurement of Respiratory Exchange Ratio and D) Metabolic Rate, measured in CLAMS cages over three consecutive days in IgMi and Ctrl animals, *n*=2-4 per group and representative of two independent experiments. All data shown as +/- SEM unless otherwise indicated, * p< 0.05, ** p< 0.01, *** p< 0.001, **** p< 0.0001.

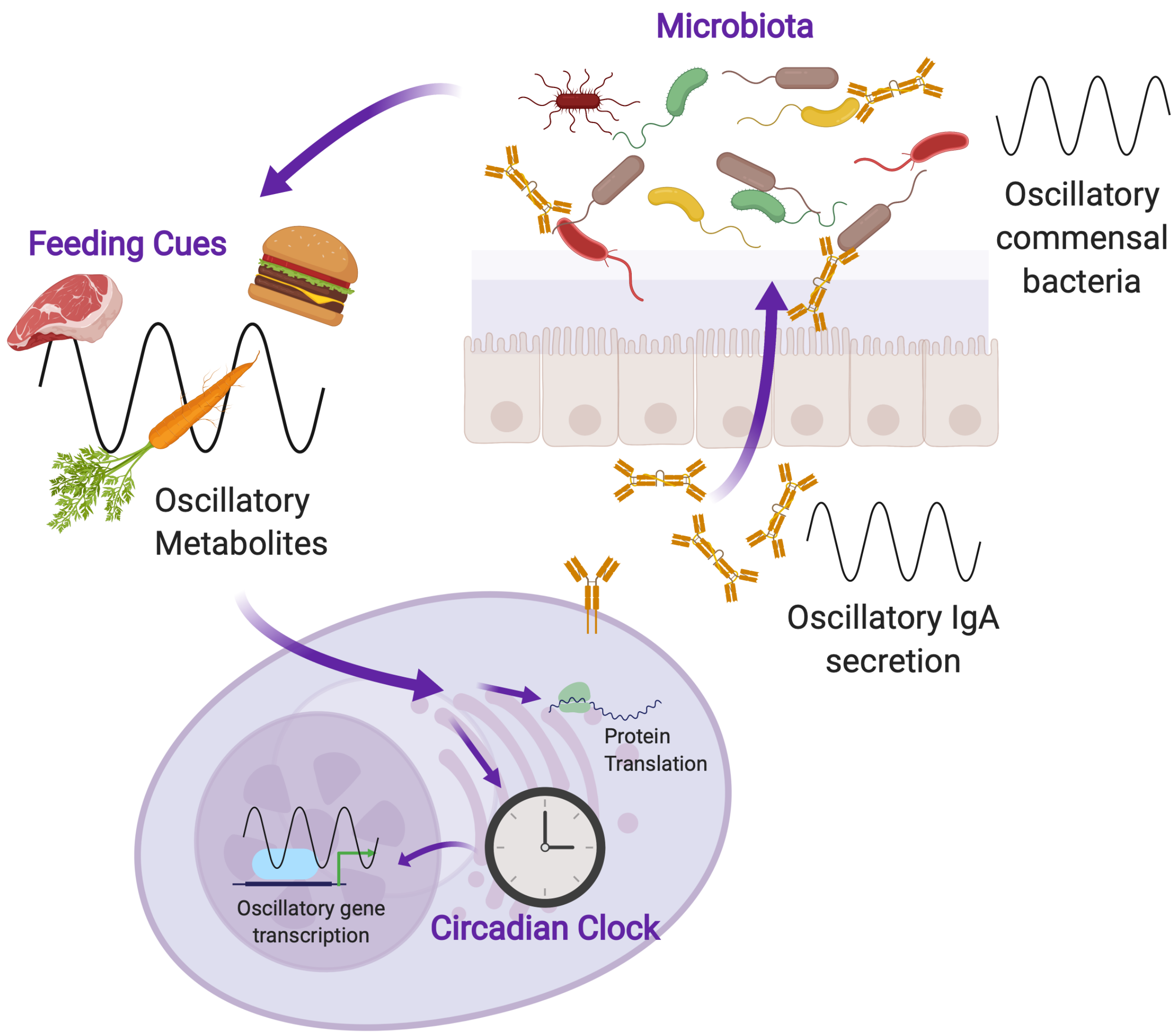


**Fig. S8. Schematic summary of circadian IgA regulation of rhythmic commensal microbiota mutualism.** Here we demonstrate diurnal oscllations in the secretion of IgA by small intestinal plasma cells. Oscillations in IgA were in part regulated by cell-intrinsic circadian clock machinery in addition to feeding-associated nutritional and metabolic cues. Oscillations in IgA act in part to regulate concurrent oscillations in the composition and relative abundance of commensal bacteria within the intestinal-resident microbiota and to influence the activity and function of gut microbes. In the absence of IgA we demonstrate perturbations in the normal circadian regulation and availability of luminal metabolites, such as glucose, which correlate with systemic uptake of glucose by the host. These data suggest that there may be a complex and reciprocal relationship between the regulation of diurnal oscillations in both IgA and microbiota that are influenced by the diet and the subsequent metabolism of nutrients by the microbiota. Image generated with BioRender.
